# Reprogramming of Osimertinib-Resistant EGFR-mutant NSCLC: The Pyruvate-Acetaldehyde-Acetate Pathway As a Key Driver of Resistance

**DOI:** 10.1101/2025.06.27.661913

**Authors:** Giorgia Maroni, Eva Cabrera San Millan, Raffaella Mercatelli, Alice Chiodi, Beatrice Campanella, Massimo Onor, Letizia Modeo, Giulia Braccini, Giovanni Cercignani, Azhar Ali, Ettore Mosca, Emilia Bramanti, Elena Levantini

## Abstract

Osimertinib (Osi) resistance remains a significant challenge in EGFR mutant non-small-cell lung cancer (NSCLC). This study investigates the metabolic reprogramming associated with Osi resistance, identifying key metabolic vulnerabilities that may be targeted for therapeutic intervention. Employing the EGFR-mutant H1975 parental (Par) cell line and its Osi-resistant (OsiR) counterpart, we integrated transcriptomics, metabolomics, nuclear and mitochondrial genomics, functional assays and bioanalytical techniques, as well as advanced 3D imaging to comprehensively define the resistant phenotype.

We found that OsiR cells exhibit mitochondrial dysfunction, including impaired oxidative phosphorylation (OXPHOS), mitochondrial DNA mutations, and altered mitochondrial gene expression. To describe this systems-level characterization, we introduce the concept of *mitochondromics*, a comprehensive profiling of mitochondrial genomic, transcriptomic, structural, and functional changes contributing to therapeutic resistance.

Metabolomic profiling revealed a significant accumulation of glycolytic intermediates (lactate, pyruvate, acetate, and acetaldehyde) in the extracellular medium, indicating a shift toward glycolysis and activation of alternative metabolic pathways, including the Warburg effect. Notably, we identified the pyruvate-acetaldehyde-acetate (PAA) pathway as a functionally repurposed metabolic route that facilitates NADPH production, which is critical for antioxidant defense and anabolic processes in OsiR cells. Additionally, although the pentose phosphate pathway (PPP) is not the primary source of NADPH in OsiR cells, it plays a supporting role in biosynthesis, contributing to the production of amino acids, nucleotides, and vitamins. Altered expression of enzymes involved in glycolysis, the TCA cycle, and both oxidative and non-oxidative arms of the PPP further supports an adaptive metabolic network promoting cell growth and resistance to Osi.

This study reveals a complex metabolic reprogramming in Osi-resistant EGFR-mutant NSCLC, where a newly identified role for the PAA pathway, alongside integrated mitochondromic alterations emerges as key driver of resistance. These insights uncover potential metabolic vulnerabilities of Osi-resistant tumors and provide a foundation for developing therapeutic strategies to counteract resistance and improve osimertinib efficacy. Targeting these metabolic pathways may offer promising avenues for overcoming resistance in clinical settings.

## INTRODUCTION

Resistance to tyrosine kinase inhibitors (TKIs) such as osimertinib represents a significant clinical challenge in treating EGFR-mutant NSCLC. The mechanisms driving osimertinib resistance are complex and multifactorial, involving both EGFR-dependent and -independent pathways (1). These include secondary mutations, activation of alternative signaling cascades, alterations in downstream effectors, and phenotypic adaptations within both cancer cells and the tumor microenvironment (TME).

Our previous work has uncovered two metabolic alterations in TKI-resistant EGFR-mutated NSCLC. First, we found that the palmitoylation of EGFR, regulated by Fatty Acid Synthase (FASN), influences EGFR stability and can be inhibited by Orlistat, partially mitigating resistance (2). Second, we showed the interaction between Caveolin-1 and the glucose transporter GLUT3 enhances glucose uptake, a process that can be disrupted by Atorvastatin, impacting resistance mechanisms (3). These findings motivated our current investigation into metabolic vulnerabilities that may be exploited to overcome TKI resistance in EGFR-mutated NSCLC.

Abnormal cellular metabolism is a hallmark of cancer, supporting tumor initiation, progression, and therapeutic resistance. Despite its importance, the mechanisms underlying this metabolic rewiring remain elusive. Cancer cells, including those in NSCLC, typically transition from mitochondrial OXPHOS to increased aerobic glycolysis, a phenomenon known as the Warburg effect. This metabolic shift supports cell survival and contributes to drug resistance (4-6). However, the role of mitochondria, central players in energy production, apoptosis regulation, and metabolic signaling, remains underexplored in the context of therapeutic resistance to TKI, such as osimertinib.

Acetate metabolism plays a critical yet underexplored role in this metabolic shift. Since the 1950s, it has been known that tumor cells preferentially utilize pyruvate or acetate for biosynthetic needs (7, 8). Under hypoxia, cancer cells often rely on alternative carbon sources like acetate to produce acetyl-CoA, highlighting the role of mitochondrial function in sustaining tumor growth and therapeutic resistance (9, 10). The ability of cancer cells to balance intracellular levels of acetyl-CoA and acetate is especially critical under nutrient-limiting conditions, such as when glucose is scarse.

Despite these observations, the interconnections between acetate metabolism, glycolysis, OXPHOS and mitochondrial dynamics in the context of TKI resistance remain fragmented. This study seeks this gap in knowledge by comprehensively investigating the role of acetate metabolism and mitochondrial reprogramming in osimertinib resistance in EGFR-mutant NSCLC, aiming to identify novel therapeutic vulnerabilities.

We adopted the H1975 NSCLC cell line, which harbors EGFR L858R/T790M mutations, a common resistance genotype, to investigate metabolic alterations associated with osimertinib resistance. Our integrated approach, combining transcriptomics, functional assays, morphological and bioanalytical assays, as well as mitochondrial DNA profiling, enabled us to identify biomarkers of therapeutic resistance. To capture the full extent of these alterations, we propose the term *mitochondromics*: a multidimensional characterization of mitochondrial structure, function, and regulation encompassing mitochondrial genomics, transcriptomics, and metabolic output.

A central focus was the analysis of the extracellular medium (ECM), or exometabolome (i.e., extracellular metabolite profiling), using chromatography-based assays (11-14) to identify metabolites that sustain cancer cell survival and proliferation. Through this comprehensive metabolic profiling, we uncovered distinct metabolic signatures, particularly elevated levels of acetate and extracellular metabolites, driving resistance to osimertinib. Furthermore, we observed a profound shift in mitochondrial dynamics, with the non-oxPPP playing a central role in providing reducing equivalents and supporting biosynthesis. Notably, we identified the pyruvate-acetaldehyde-acetate (PAA) pathway as a key contributor to resistance, driving the generation of critical metabolites, such as NADPH, which is essential for reductive biosynthesis, and acetyl-CoA, which supports lipid biosynthesis. This pathway serves as a central hub in the metabolic reprogramming that supports resistance development.

## SIGNIFICANCE

By integrating metabolomic, transcriptomic, genomic, functional and bioanalytical approaches, this study provides a comprehensive understanding of the complex metabolic network driving osimertinib resistance in EGFR-mutant NSCLC. We identify the **pyruvate-acetaldehyde-acetate (PAA) pathway** as a previously unrecognized metabolic route that facilitates NADPH production, a critical factor in sustaining cell proliferation and drug resistance. We further connect this pathway to essential biosynthetic processes and mitochondrial reprogramming. Through the lens of *mitochondromics*, our findings highlight the dual role of mitochondria as both sensors and effectors of metabolic adaptation in the resistant phenotype. These insights uncover new metabolic vulnerabilities in EGFR-mutant tumors and suggest promising strategies to enhance osimertinib efficacy by overcoming resistance mechanisms.

## 2. RESULTS

### 2.1 RNA sequencing reveals the glycolytic adaptation of Osimertinib-resistant EGFR mutant NSCLC cells

To investigate the mechanisms underlying osimertinib resistance, we performed RNA sequencing (RNAseq) to analyze transcriptional changes in Par H1975 NSCLC cells as they adapted to the drug and developed resistance. Differential gene expression (DGE) analysis identified a broad set of dysregulated genes in OsiR cells, which were selected based on a threshold of a two-fold change (LogFC ≥ 0.25, FDR ≤ 0.05) in gene expression.

Enrichment analysis using EnrichR (**Figure 1a**) revealed the significant involvement of mitochondrial processes in OsiR cells, including *Mitochondrial Translation, Mitochondrial Gene Expression*, and *Cellular Response to Oxidative Stress*, within the Gene Ontology (GO) Biological Process (BP) category (adjusted p-value <0.05). In line with this, mitochondrial-associated components, including *Mitochondrial Ribosomes, Mitochondrial Membrane*, and *Mitochondrial Inner Membrane* were prominently represented in the GO Cell Compartment (CC) category, emphasizing the pivotal role of mitochondria in resistance development. These findings support the hypothesis that mitochondrial adaptation is central to cellular resistance mechanisms that emerge under Osi-induced stress.

**Figure 1.**
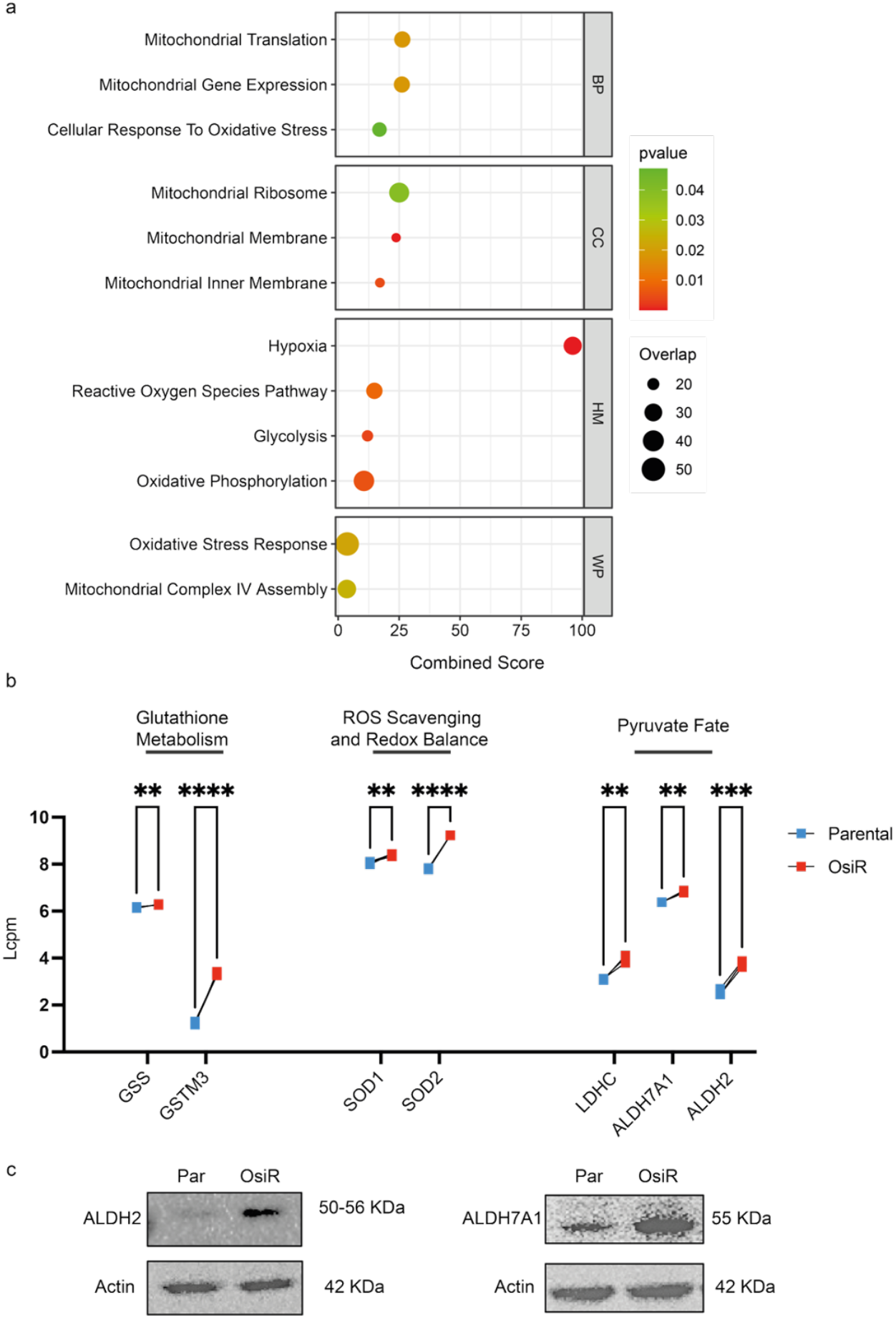
Transcriptomic analysis highlights metabolic adaptation in Osimertinib-resistant EGFR mutant NSCLC cells. **(a)** Pathway enrichment analysis of differentially expressed genes in OsiR versus Par H1975 cells using EnrichR. Upregulated pathways are shown across Gene Ontology (GO) categories, Biological Processes (BP) and Cellular Components (CC), as well as Hallmark (HM) and WikiPathways (WP). Dot size indicates gene overlap; color represents adjusted p-value. **(b)** RNA-seq analysis showing increased expression of genes involved in glutathione metabolism, ROS scavenging and redox balance, as well as pyruvate fate in H1975 Par (blue) and OsiR (red) cells. Statistical significance is indicated (**p<0.0021 to ****p<0.0002). **(c)** Western blot analyses of H1975 Par and OsiR cells. Protein lysates were immunoblotted with antibodies against ALDH2 and ALDH7A1. β-actin was used as a loading control. Expected molecular weights (kDa) are indicated.

Consistent with this, HallMark (HM) pathway analysis identified enrichment in *Hypoxia* and *Reactive Oxygen Species (ROS) Pathway*, reinforcing the importance of redox regulation in the resistant phenotype. Additionally, HM analysis highlighted *Glycolysis* and *Oxidative Phosphorylation*, suggesting a dynamic shift in cellular energy metabolism. The enrichment in *Glycolysis* reflects a well-documented adaptive strategy employed by cancer cells facing therapeutic pressure, indicating that OsiR cells undergo metabolic reprogramming toward glycolysis to support survival despite drug treatment.

Further support for this metabolic remodeling was provided by WikiPathway (WP) analysis, which confirmed the involvement of *Oxidative Stress Response* and *Mitochondrial Complex IV Assembly*, reinforcing the link between mitochondrial homeostasis and acquired drug resistance.

In line with these findings, the observed significant upregulation of genes involved in ROS generation across multiple platforms, including GO-BP, HM and WP, indicative of heightened oxidative stress in OsiR cells. This stress is likely driven by mitochondrial dysfunction and intensified metabolic activity. The balance between ROS production and scavenging is crucial for tumor cell survival; while excessive ROS can induce cellular damage, moderate ROS levels can promote survival and facilitate adaptation to stress.

Supporting this, transcriptomic data revealed the upregulation of genes involved in glutathione metabolism, such as *glutathione synthetase (GSS)* and *glutathione S-transferase Mu 3 (GSTM3)*, alongside ROS-scavenging genes like *superoxide dismutase 1 (SOD1)* and *superoxide dismutase 2 (SOD2)* (**Figure 1b**). These changes suggest an enhanced detoxification response in OsiR cells, enabling them to better manage oxidative stress.

In addition to ROS regulation, our transcriptomic analysis also showed increased expression of metabolic enzymes associated with cancer progression (15-18), including *Lactate Dehydrogenase C (LDHC), Aldehyde Dehydrogenase 2 (ALDH2)* and *Aldehyde Dehydrogenase 7 Family Member A1 (ALDH7A1)* (**Figure 1b**). Western Blot analysis confirmed the upregulation of ALDH2 and ALDH7A1 at the protein level (**Figure 1c**). These enzymes not only participate in aldehyde detoxification but also contribute to metabolic resilience under stress, reinforcing the model in which OsiR cells engage in extensive metabolic rewiring.

This rewiring includes a strategic realignment of mitochondrial function, what could be defined as a *mitochondriomic shift*, geared toward maintaining redox balance and sustaining survival under therapeutic pressure.

### 2.2 Distinct Mitochondrial Morphological Alterations in Osimertinib-resistant EGFR mutant NSCLC cells

To investigate the link between metabolic remodeling and mitochondrial architecture, we employed 3D mitochondrial reconstruction by confocal microscopy on Par and OsiR cells. OsiR cells exhibited a fragmented, toroidal mitochondrial morphology with reduced size, indicative of a disrupted mitochondrial network. In contrast, Par cells displayed elongated, interconnected mitochondria characteristic of a healthy, fused network (**Figure 2a**). These observations align with the idea that mitochondrial fragmentation is a hallmark of cellular stress and drug resistance.

**Figure 2.**
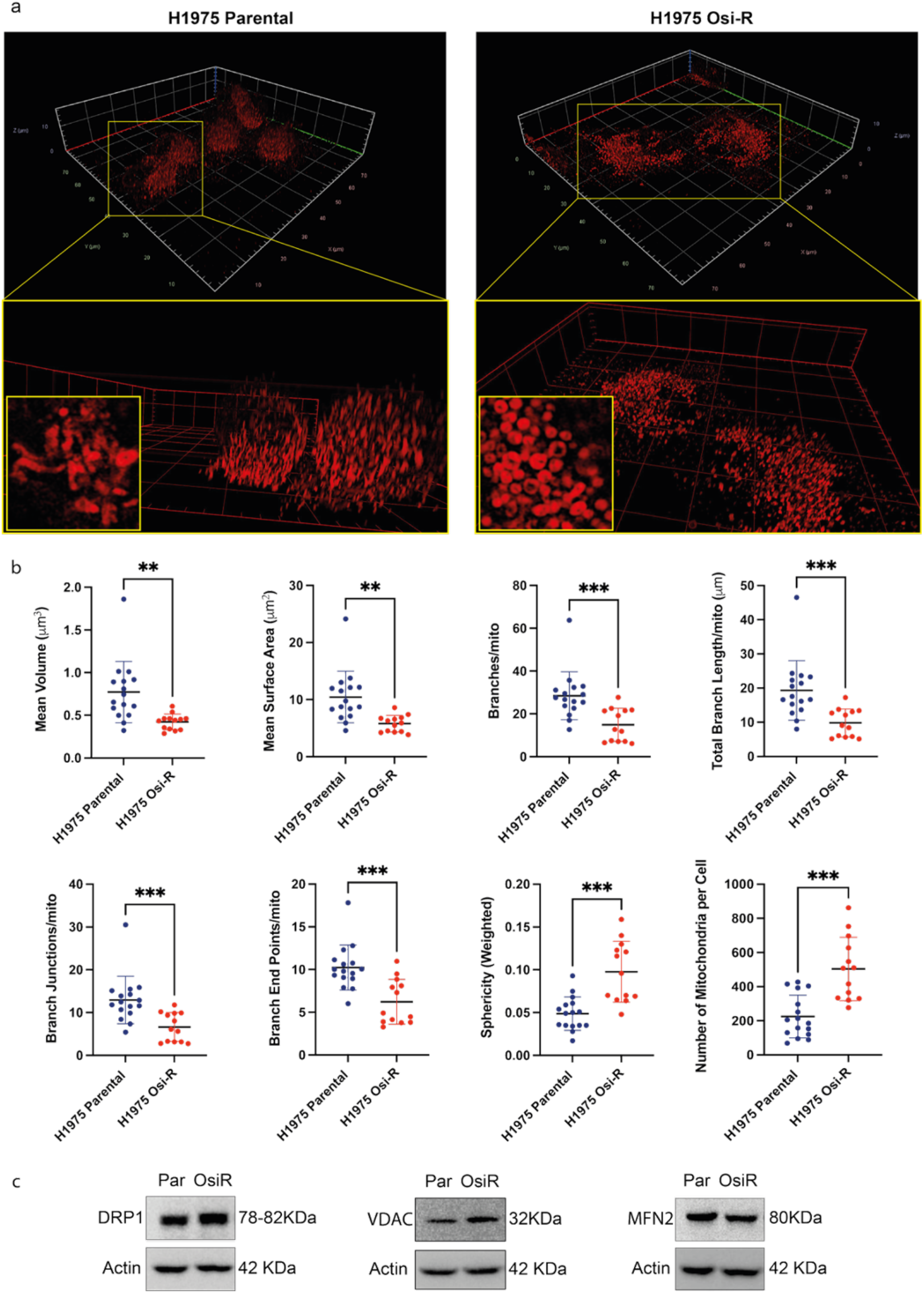
Mitochondrial imaging and protein analysis highlight structural and dynamic remodeling in Osimertinib-resistant EGFR mutant NSCLC cells. **(a)** Leica Airyscan confocal microscopy 3D reconstruction of mitochondria in H1975 cells. MITO-ID staining (red channel) of mitochondria in Par H1975 cells (left panel) and in OsiR H1975 cells (right panel). Insets show zoomed-in views of mitochondrial architecture. **(b)** Quantitative 3D morphometric analysis of mitochondria performed using the Mitochondria Analyzer plugin in Fiji. Error bars represent standard deviation (SD); p-values are indicated (**p<0.0021 to ***p<0.0002). **(c)** Western blot analyses of H1975 Par and OsiR cells. Protein lysates were immunoblotted with antibodies against DRP1, VDAC and MFN2. β-actin was used as a loading control. Expected molecular weight (kDa) is indicated.

Quantitative 3D confocal analysis confirmed these structural differences. OsiR cells showed significant reductions in mitochondrial volume, surface area, total branch number, branch length, junctions, and endpoints per mitochondrion. Additionally, sphericity was markedly increased, and mitochondrial number per cell was higher (**Figure 2b**), supporting a shift toward a fragmented and stress-responsive mitochondrial phenotype under osimertinib pressure.

Western blot analysis further validated this shift in mitochondrial dynamics. OsiR cells exhibited increased expression of Dynamin-Related Protein 1 (DRP1), a key driver of mitochondrial fission (**Figure 2c**). In contrast, Mitofusin 2 (MFN2), a central regulator of mitochondrial fusion, was more abundant in Par cells (**Figure 2c**). Additionally, levels of Voltage-Dependent Anion Channel (VDAC), a marker of outer mitochondrial membrane integrity, were observed in OsiR cells (**Figure 2c**), reinforcing the notion of altered mitochondrial dynamics in resistant cells.

This morphological reprogramming further underscores the central role of mitochondria in orchestrating adaptive responses to therapeutic stress.

### 2.3 Mitochondrial and Nuclear Genome mutations as Bioenergetic Adaptations in Osimertinib Resistance

To investigate the bioenergetic adaptations underlying osimertinib resistance, we performed targeted mitochondrial DNA sequencing (mtDNA-seq) alongside whole genome sequencing (WGS) of nuclear DNA in Par and OsiR cells. This dual-genome approach uncovered a coordinated pattern of mutational remodeling affecting both mitochondrial and nuclear-encoded OXPHOS genes, suggesting a concerted genomic response to drug pressure.

### Mitochondrial Genome Remodeling in OsiR cells

Mitochondrial genome sequencing identified a total of 171 single-nucleotide variations (SNVs) scattered throughout the mitochondrial DNA, predominantly at low variant allele frequency (AF) **(Suppl. Table 1 and Suppl. Data 1)**. Most variants were predicted to have moderate impact, with a smaller subset (7/171) classified as high-impact category (e.g., stop-gained or start-lost mutations).

When comparing Par and OsiR cells, we found that 128 mutations (74.85%) were already present in Par cells (113 were Par-specific, and 15 shared with OsiR but showed higher AF in Par cells). In contrast, 43 mutations (25.15%) were associated with resistance: 38 emerged specifically in OsiR cells, and 5 were present in both but enriched in OsiR cells. By focusing on condition-specific mutations, i.e. those completely absent in the other condition (AF = 0), we identified 113 as Par-specific (66.08%), 38 as OsiR-specific (22.22%), and 20 shared between both (11.70) **(Suppl. Table 1)**.

These data suggest that resistance acquisition involves both purging of pre-existing mutations and gaining of novel ones, reflecting dynamic mitochondrial genome remodeling. Notably, the turnover was particularly prominent within ETC-encoded subunits. Complex I (C-I) was the most affected, with 93 variants overall: 56 Par-specific (fully lost in OsiR), 14 shared (3 enriched in OsiR; 11 in Par), and 23 newly-acquired in OsiR cells. This represents an ∼80% purging of Par-derived mutations. Complex III (C-III) showed 10 OsiR-specific variants and 2 shared ones, with divergent retention in the two cell types. Complex IV (C-IV) underwent a drastic 94% reduction in Par-specific mutations (47 in total), retaining 3 shared variants (all enriched in Par), and acquiring only 2 OsiR-specific ones. Complex V (C-V) also lost 91% of its Par-derived mutations (10 total), while gaining 3 new OsiR-specific variants and retaining only one shared mutation with increased AF in OsiR cells (**Suppl. Data 1**). Collectively, these patterns suggest a non-random, selective editing of the mitochondrial genome during resistance development. The mutational landscape appears to shift toward a more functionally favorable configuration, potentially optimizing mitochondrial performance under drug pressure.

### Nuclear Genome Alterations in OXPHOS genes

WGS identified 121 mutations in nuclear-encoded genes involved in OXPHOS, primarily consisting of SNVs and a smaller number of short insertions and deletions (indels) (**Suppl. Table 1 and Suppl. Data 2**). All variants were predicted to be modifiers, suggesting they may influence gene expression or protein function without causing complete loss of function.

Notably, 87 of these mutations (71.9%) were more abundant (displaying a higher allele frequency) in Par cells, while only 34 (28.1%) were enriched in OsiR cells. This pattern of mutual loss in OsiR cells suggests genetic restructuring during the acquisition of drug resistance (**Suppl. Table 1**).

When mutations were mapped across ETC complexes, distinct complex-specific patterns emerged. C-I carried 32 mutations evenly split between variants enriched in Par and OsiR cells (15 and 17, respectively). C-IV showed a similarly balanced distribution, with 14 Par- and 12 OsiR-enriched ones. In contrast, C-III displayed complete asymmetry were all 6 detected mutations were enriched in Par cells and diminished in OsiR cells, implying selective depletion. C-II, fully encoded by the nuclear genome, carried only 3 mutations in total (2 enriched in Par, 1 in OsiR).

Remarkably, C-V showed the strongest asymmetry, with 49 of 52 variants diminished in OsiR cells, and only 3 showing increased AF, suggesting this complex may be under heightened selective pressure in the resistant state.

The indel landscape followed a similar trend. Multiple deletions and insertions, such as a 17-nt deletion and a 4-nt insertion in C-I, a 14-nt insertion and two 2-nt substitutions in C-IV, and several insertions of varying length (2-, 3-, 11-, and 18-nt) in C-V, were all reduced in OsiR cells (**Suppl. Data 2**). This consistent reduction further supports the concept of a selective pruning mechanism during resistance evolution.

Together, these findings indicate that osimertinib resistance is associated with active remodeling of the nuclear genome within OXPHOS genes. The preferential retention of certain mutations and the elimination of others suggest a process of genetic optimization, where deleterious or unnecessary variants are eliminated.

To further refine the landscape of OXPHOS remodeling, we next analyzed gene-specific alterations within each ETC complex, integrating mitochondrial- and nuclear-encoded components to identify key mutational targets.

**Supplementary Table 1.** Mutations that are specific in Par or have higher AF in Par are displayed in blue. Mutations that are specific in OsiR or have higher AF in OsiR are displayed in Red. Grey background applies to mtDNAseq while light blue background applies to WGS.

|  |  |  | shared | shared |  |  |  |  |
| --- | --- | --- | --- | --- | --- | --- | --- | --- |
|  | Par- Specific | OsiR-Specific | Par>OsiR | OsiR>Par | TOT | Par>OsiR | OsiR>Par | TOT |
| Complex I | 56 | 23 | 11 | 3 | 93 | 14 | 17 | 31 |
| Complex I;Complex IV |  |  |  |  |  | 1 |  | 1 |
| Complex II |  |  |  |  |  | 2 | 1 | 3 |
| Complex III |  | 10 | 1 | 1 | 12 | 6 |  | 6 |
| Complex IV | 47 | 2 | 3 |  | 52 | 14 | 11 | 25 |
| Complex IV;Respirasome assembly |  |  |  |  |  |  | 1 | 1 |
| Complex V | 10 | 3 |  | 1 | 14 | 49 | 3 | 52 |
| Cytochrome C |  |  |  |  |  | 1 |  | 1 |
| Respirasome assembly |  |  |  |  |  |  | 1 | 1 |
| Total (by condition) | 113 | 38 | 15 | 5 | 171 | 87 | 34 | 121 |
| Sequencing (mtDNA-seq or WGS) | 171 |  |  |  | 171 | 121 |  | 121 |

All seven mtDNA-encoded subunits showed dynamic changes in their mutational profiles. In OsiR cells, MT-ND1 and MT-ND2 lost most of their Par-specific mutations (14 and 34, respectively), retaining only a few (6 and 5) at reduced AFs, while acquiring a small number of OsiR-specific mutations (2 and 4, respectively). MT-ND5 followed a similar trajectory, eliminating all its 7 Par mutations and gaining three new ones during resistance. In contrast, MT-ND3 and MT-ND4 showed net gains of five new variants each, with MT-ND4 losing only a single Par-specific mutation. Notably, some mutations were not only retained but expanded in OsiR cells, such as one in MT-ND3 and one in MT-ND4 that nearly tripled in AF. MT-ND4L and MT-ND6 similarly expanded their mutation load in the resistant state, the former acquiring 4 new variants, and the latter amplifying a pre-existing Par-derived mutation. Together, these patterns highlight a dynamic and selective remodeling of mtDNA during resistance acquisition.

Nuclear-encoded subunits of Complex I displayed a similarly complex and coordinated remodeling pattern, with 32 distinct variants identified across 21 subunits (**Figure 3b**). A subset of these variants showed clear positive selection, with significantly increased AFs in OsiR cells. This group included assembly factors and accessory or catalytic core subunits such as NDUFAF6 (three variants, one up to 15.5-fold increase), NDUFB1 (8.6-fold), NDUFA7 (7.9-fold), and NDUFS2 (6.7-fold). Other positively selected subunits included NDUFA6, NUBPL (two variants), LYRM2, NDUFA12, NDUFB9 and NDUFAF2, with increases ranging from 6.4- to 4-fold, suggesting that remodeling of these elements may enhance Complex I stability or activity under drug pressure. Moderate enrichment (1.4–2.75-fold) was observed in NDUFA9, NDUFS4, and NDUFB2 (two variants), which support structural and electron transfer functions. In contrast, several mutations were negatively selected during resistance. These included variants in NDUFAF2 (two variants, one up to an 8.7-fold decrease), NDUFA5 (6.3-fold), NDUFS7, NDUFAF6, NDUFA10, NUBPL (three variants), NDUFC1, NDUFV2, NDUFAB1, NDUFA8 (two variants), NDUFAF2, and COA1 (a Complex IV assembly factor), with reductions ranging from 4.73- to 1.9-fold. Mixed mutational patterns were observed in NDUFAF6, NDUFAF2, and NUBPL, each carrying both positively and negatively selected variants, suggesting that Complex I remodeling is not a simple binary switch but involves a finely tuned, selective adaptation of both mitochondrial and nuclear subunits.

**Figure 3.**
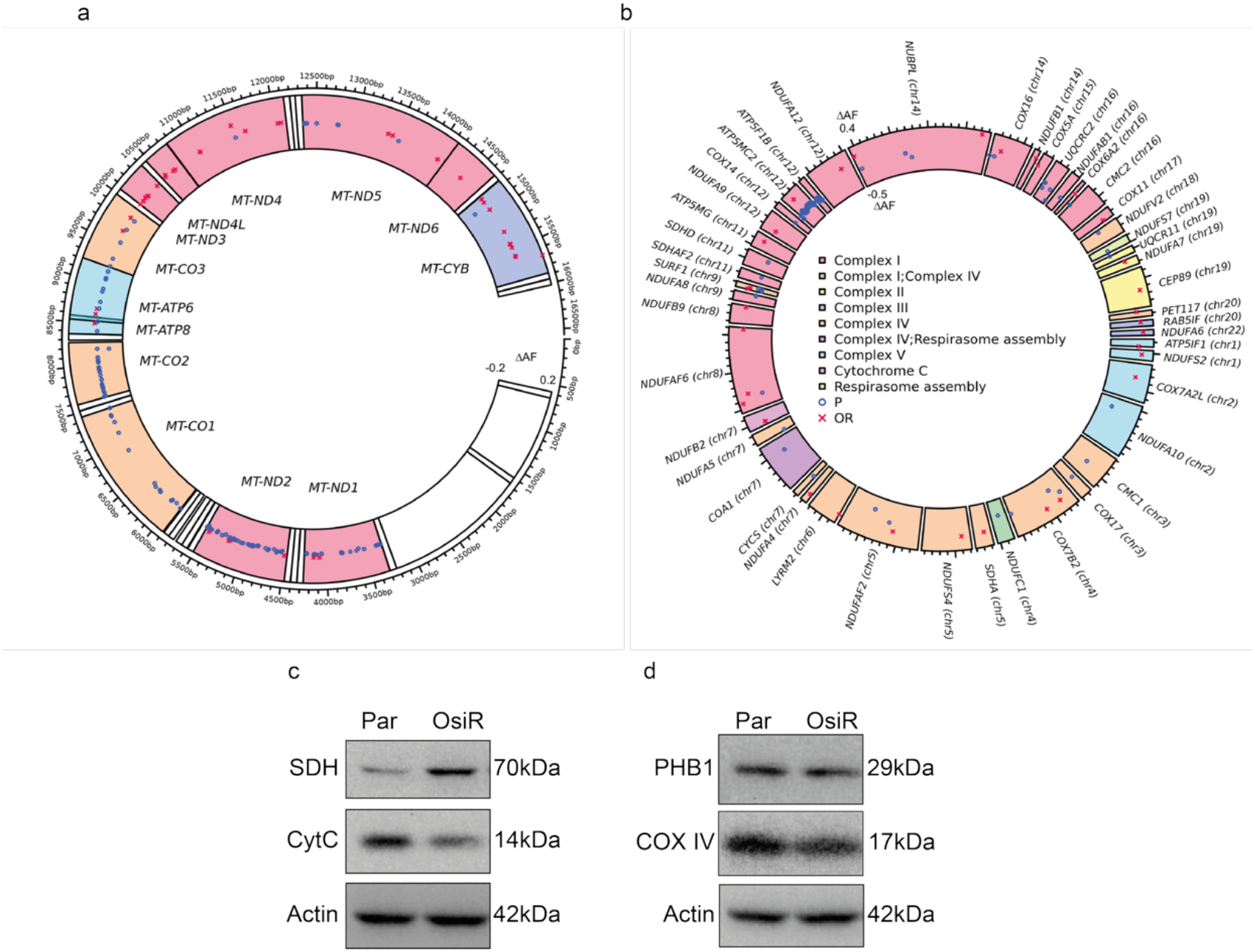
Mitochondrial and nuclear genome remodeling in Osimertinib-resistant EGFR mutant NSCLC cells. **(a)** Mitochondrial and **(b)** nuclear genome mutations in H1975 Par versus OsiR cells, mapped by genomic location (across the circle) and difference in Allele Frequency (ΔAF, radial axis). Blue circles indicate mutations present in Par cells that were lost in OsiR cells (ΔAF < 0); red crosses represent mutations gained in OsiR cells (ΔAF > 0). Mitochondrial genes are color-coded by respiratory complex. **(c-d)** Western blot analyses of H1975 Par and OsiR cells. Protein lysates were immunoblotted with antibodies against SDH, CytC, PHB1 and COX IV. β-actin was used as a loading control. Expected molecular weights (kDa) are indicated.

### Remodeling of Complex II

Unlike Complex I, Complex II (succinate dehydrogenase, SDH) is entirely encoded by nuclear DNA and is composed of SDHA and SDHB (catalytic core), together with SDHC and SDHD (anchoring the complex to the membrane). C-II displayed minimal mutational changes. WGS identified only three SNVs across Par and OsiR cells, with two enriched in Par and one in OsiR cells (**Figure 3b**). The only variant showing a notable AF increase (7.63-fold) in OsiR cells was located in *SDHA*, the catalytic subunit of SDH.

Overall, these findings suggest that C-II remains largely insulated from mutational remodeling, even under strong drug selection pressure. However, despite minimal mutational changes, the increased expression of SDH in OsiR cells (**Figure 3c**) suggests the presence of non-genetic compensatory mechanisms, possibly at the post-transcriptional or translational level, aimed at reinforcing OXPHOS output in response to its dysregulation under drug pressure.

### Remodeling of Complex III

Complex III (cC-III, cytochrome bc_1_ complex) is a multi-heteromeric enzyme composed of 11 different subunits per monomer, that assemble into a symmetric homodimer (CIII_2_), which constitutes the functionally active enzyme complex. All subunits are encoded by nuclear genes, except for one (MT-CYB), which is encoded by mtDNA(19). MT-CYB is central to electron transfer, and mtDNA-seq revealed 12 SNVs, of which 10 (83.3%) were OsiR-specific and only two were shared with Par cells (**Figure 3a**). WGS also identified six nuclear-encoded variants, predominantly affecting **UQCRC2** (five variants) and **UQCR11** (one variant), which are involved in electron transfer and C-III assembly. These nuclear variants were consistently reduced in OsiR cells, with AFs decreasing by up to 9.63-fold (**Figure 3b**).

This genomic remodeling was accompanied by reduced MT-CYB protein levels in OsiR cells (**Figure 3c**), as well as decreased levels of Prohibitin 1 (PHB1), a critical scaffold protein for C-III assembly and stability (**Figure 3d**). Together, these findings suggest that C-III destabilization is a hallmark of the resistant mitochondrial phenotype.

### Cytochrome c Variant

In addition, *CYCS*, which encodes cytochrome c, the mobile electron shuttle between C-III and C-IV, carried a single nuclear variant showing a 6.33-fold reduction in OsiR cells (**Figure 3b; Suppl. Table 1 and Suppl. Data 2**), potentially impacting electron flow efficiency.

### Remodeling of Complex IV

Complex IV (cytochrome c oxidase) is the terminal enzyme of the mitochondrial OXPHOS system, catalyzing electron transfer from cytochrome c to oxygen, the final electron acceptor, while concurrently pumping protons across the inner mitochondrial membrane. This process contributes to the proton motive force essential for ATP synthesis via Complex V(20). C-IV consists of 14 subunits: three encoded by mtDNA (MT-CO1, MT-CO2, and MT-CO3), which form the catalytic core, and 11 nuclear-encoded structural and regulatory subunits.

mtDNA-seq identified 52 mitochondrial variants affecting C-IV genes. Of these, 47 (95.2%) were completely lost during resistance acquisition, with only three variants (4.8%) retained at reduced AFs, one in MT-CO1 and two in MT-CO2 (**Figure 3a, Suppl. Table 1 and Suppl. Data 1**), suggesting selective pruning of parental mutations under therapeutic pressure. Additionally, two novel mutations emerged exclusively in OsiR cells, indicative of clonal diversification during adaptation.

Nuclear sequencing uncovered 27 additional variants in C-IV-related genes, with a clear redistribution between Par and OsiR cells: 15 variants were enriched in Par and 12 in OsiR cells (**Figure 3b, Suppl. Table 1**). These included mutations in both structural subunits and biogenesis regulators. Among them NDUFA4, previously misclassified as a Complex I subunit but now recognized as a *bone fide* C-IV component involved in C-IV function and biogenesis, exhibited a 7.53-fold increase in AF in OsiR cells (**Suppl. Data 2**). SURF1, a key assembly factor, emerged as a mutational hotspot with six distinct variants: two enriched in OsiR cells (4.5- and 6.1-fold), and four reduced (by 2.5-7.6 fold), including the only insertion identified among C-IV-related genes. Other nuclear-encoded subunits also exhibited notable changes: COX14 and COX17 were consistently reduced (up to ∼5-fold), whereas COX11, COX5A, COX6A2, COX7A2L, PET117 and CEP89 displayed AF increases ranging from 2.9- to 5.5-fold in OsiR cells. COX16 and COX7B2 showed mixed dynamics, displaying both enriched, reduced, and also unchanged variants. By contrast, subunits involved in stabilizing and assembling the complex, including CMC1, CMC2 and COA1 were uniformly decreased, with COA1 showing a 50% reduction in AF. Notably, both COA1 and COX7A2L serve dual roles, participating not only in C-IV biogenesis but also in Complex I function and in mitochondrial supercomplex (respirasome) formation, underscoring the interconnectedness of respiratory chain regulation.

Transcriptomic data supported this genomic remodeling. WikiPathway enrichment analysis revealed significant upregulation of the *Mitochondrial Complex IV Assembly* pathway in OsiR cells (**Figure 1a**), indicating transcriptional reinforcement of C-IV biogenesis and structural optimization. Despite no mutations in COXIV, its protein levels were reduced in OsiR cells (**Fig. 3d**) without corresponding changes in mRNA expression (data not shown), pointing to translational or post-translational modulation, underlining the multilayered nature of mitochondrial adaptation.

Together, these findings reveal extensive restructuring of Complex IV during resistance acquisition. Coordinated gains and losses of mitochondrial and nuclear variants, especially in structural subunits and biogenesis factors, illustrate selective optimization of ETC components. These adaptations likely contribute to the altered respiratory phenotype observed in OsiR cells, positioning C-IV as a key node in dynamic respiratory rewiring.

### Remodeling of Complex V

Complex V (ATP synthase) is the final enzyme of the OXPHOS system, responsible for ATP production, leveraging the proton gradient generated by upstream ETC complexes (I-IV). C-V consists of 29 subunits, two of which (MT-ATP6 and MT-ATP8) are encoded by the mitochondrial genome and are essential for proton translocation within the membrane-embedded F_0_ domain(21).

mtDNA-seq revealed distinct alterations in C-V composition between Par and OsiR cells (**Figure 3a, Suppl. Table 1 and Suppl. Data 1**), suggesting active remodeling of ATP synthase under therapeutic pressure. Specifically, three mutations (one in MT-ATP6 and two in MT-ATP8) were exclusively detected in OsiR cells, suggesting they likely emerged during drug exposure. An additional MT-ATP6 mutation already present in Par cells expanded 1.7-fold in OsiR cells (**Suppl. Data 1**). In contrast, eight MT-ATP6 and two MT-ATP8 mutations originally present in Par cells were completely lost in OsiR cells (**Suppl. Data 1**), reflecting the elimination of 80% and 50% of Par-specific variants, respectively. This pattern of selective turnover, marked by the loss of ten parental mutations and the acquisition of three novel ones, suggests a non-random selection process and supports a model of dynamic mitochondrial adaptation in response to drug pressure.

Complex V also exhibited the most pronounced asymmetry in variant distribution among OXPHOS complexes. WGS revealed a stark contrast, with 49 out of 52 variants enriched in Par cells, while only 3 were enriched in OsiR cells (**Figure 3b, Suppl. Table1 1 and Suppl. Data 2**). This skewed distribution draws attention to a targeted metabolic adaptation that converges on ATP synthase function during resistance development.

Among the nuclear-encoded subunits, both positively and negatively selected variants were observed. A few mutations were enriched in OsiR cells, including one in ATP5MC2 (11.04-fold increase), ATP5MG (9.08-fold), and ATP5IF1 (2.89-fold), suggesting they may confer adaptive advantages. Conversely, other subunits exhibited strong negative selection: ATP5F1B showed 11 distinct mutations reduced by 11.68- to 3.36-fold, and ATP5MC2 harbored a staggering total of 38 variants, decreased by 9.48- to 2.66-fold. ATP5MC2 also carried one variant specifically enriched in OsiR by 11-fold, likely reflecting distinct functional impacts depending on the specific mutation.

Although most Complex-V mutations were SNVs, ATP5MC2 also harbored four small deletions (2-18 nucleotides), all of which were significantly reduced in OsiR cells (7.25- to 3.18-fold) (**Suppl. Data 2**), possibly contributing to destabilization of the complex.

Together, these observations indicate that resistant cells undergo a coordinated metabolic rewiring characterized by reduced reliance on mitochondrial ATP production. This shift likely reflects a broader bioenergetic transition toward alternative energy sources, such as aerobic glycolysis, consistent with the Warburg effect. The selective emergence and loss of specific mitochondrial and nuclear variants suggest methodical reprogramming of mitochondrial function to maintain viability under therapeutic stress. More broadly, the mutational landscape across the OXPHOS system point to a systemic and directional reorganization shaped by selective pressures. While Complex V emerges as a central remodeling hub, Complexes I and IV also exhibit substantial variant remodeling, underscoring the metabolic plasticity of the mitochondrial network in therapy resistance.

### Respirasome Assembly Remodeling

Beyond mutations in core ETC subunits, we identified alterations in genes involved in respirasome architecture, suggesting structural reorganization of the mitochondrial ETC during resistance acquisition. Notably, *COX7A2L*, a known stabilizer of Complex III/IV supercomplexes and an accessory subunit of Complex IV (22), and *RAB5IF*, a regulator of mitochondrial membrane dynamics and cristae morphology critical for OXPHOS efficiency(23), emerged as potential contributors to ETC integrity and organization. Both genes showed increased AFs in OsiR cells (4.14-fold for COX7A2L and 8.76-fold for RAB5IF) (**Figure 3b, Suppl. Table 1 and Suppl. Data 2**), implicating auxiliary roles in supporting mitochondrial adaptability under therapeutic pressure.

These findings emphasize that mitochondrial remodeling in resistance extends beyond canonical ETC subunits, including peripheral regulators of organelle architecture and bioenergetic flexibility.

Altogether, these data reveal a sophisticated and systemic reorganization of the mitochondrial network in response to drug resistance. Remodeling of the ETC, particularly Complex V, along with the selective emergence and loss of both mitochondrial and nuclear variants, underlies an adaptive rewiring of bioenergetic pathways. This metabolic plasticity allows cancer cells to dynamically shift their energy production strategies, ensuring survival under therapeutic pressure and contributing to the persistence of drug-resistant phenotypes.

### 2.4 Impaired Mitochondrial Function in Osimertinib-resistant EGFR mutant NSCLC cells

While the specific functional consequences of individual mitochondrial variants remain to be elucidated, the consistent genomic and transcriptional alterations observed in OsiR cells suggest a coordinated rewiring of mitochondrial regulation. To determine whether these molecular changes translate into functional mitochondrial alterations, we compared the bioenergetic profiles of Par and OsiR cells.

Using the Seahorse XF Analyzer, we performed real-time metabolic flux analysis to assess oxygen consumption rates (OCRs) as a direct readout of mitochondrial respiratory activity. After measuring basal respiration, cells were sequentially treated with oligomycin (ATP synthase inhibitor) to quantify ATP-linked respiration and proton leak; FCCP (a protonophore that collapses mitochondrial membrane potential) to assess maximal respiratory capacity; and a combination of antimycin A and rotenone (complex III and I inhibitors) to measure non-mitochondrial respiration. This sequential treatment allowed quantification of key mitochondrial parameters, including basal and maximal respiration, spare respiratory capacity, ATP production, and coupling efficiency. All experiments were performed in biological triplicates (n=3).

OsiR cells displayed a bioenergetic profile indicative of mitochondrial dysfunction (**Figure 4**). Although basal OCR was comparable between Par and OsiR cells, indicating preserved baseline mitochondrial respiration, OsiR cells exhibited a significant reduction in ATP-linked OCR, reflecting impaired coupling efficiency in ATP synthesis. This deficit may result from increased proton leak or cytosolic acidification, both of which interfere with efficient OXPHOS. Indeed, proton leak was elevated in OsiR cells, suggesting instability of the inner mitochondrial membrane or dysregulated proton gradient maintenance. In addition, spare respiratory capacity was markedly reduced in OsiR cells, reflecting a compromised ability to adapt to increased energetic demands under metabolic stress.

**Figure 4.**
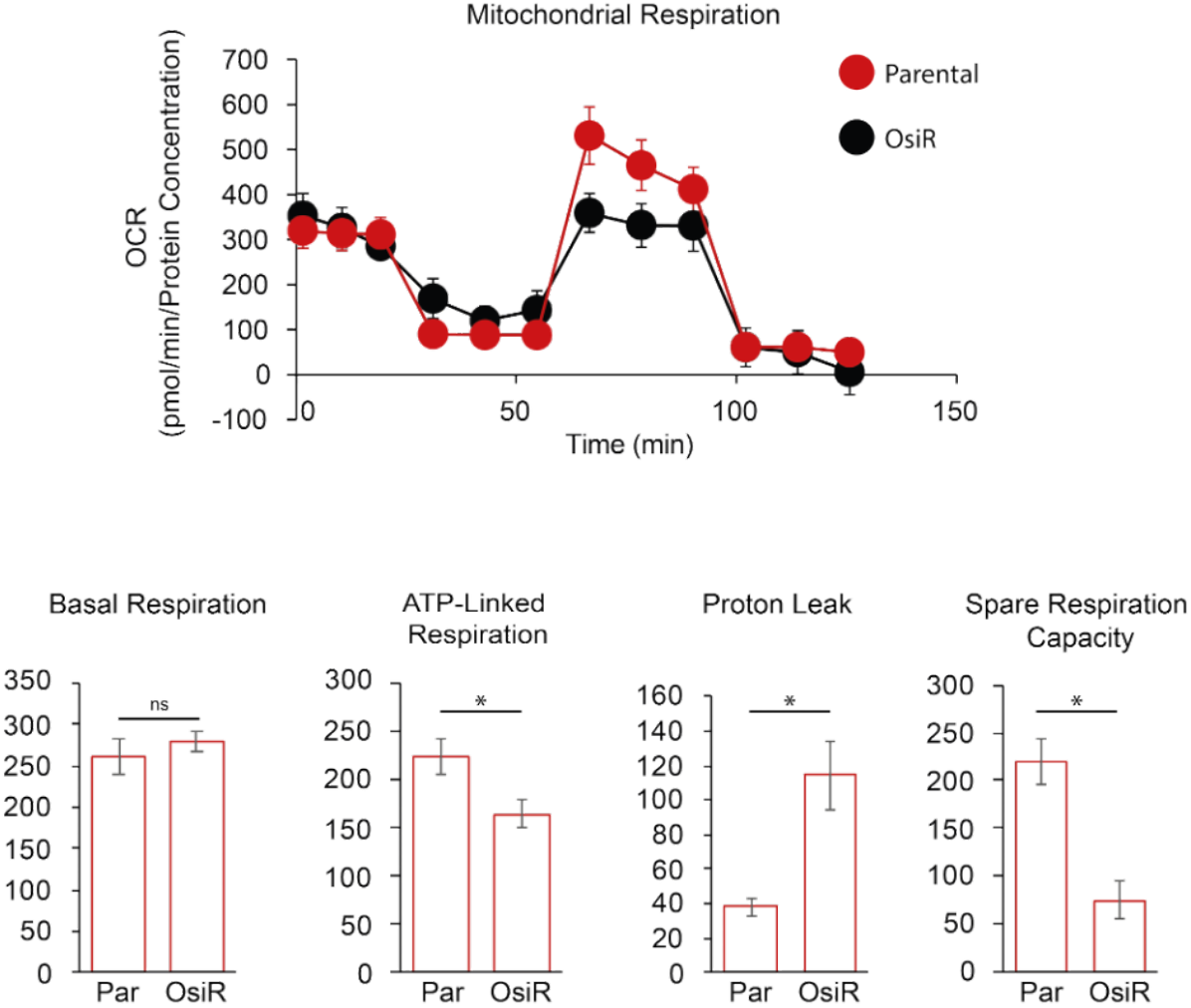
Seahorse analysis reveals impaired mitochondrial respiration and reduced oxidative capacity in Osimertinib-resistant EGFR mutant NSCLC cells. Real-time measurement of oxygen consumption rate (OCR) in H1975 Par (red) and OsiR (black) cells using the Seahorse XF Analyzer. The top graph displays dynamic OCR profiles over time. The lower panels show quantification of key mitochondrial parameters: basal respiration, ATP-linked respiration, proton leak, and spare respiratory capacity. Error bars represent standard deviation (SD); *p*-values are indicated (ns, *p* > 0.05; * *p* < 0.05).

Together, these data demonstrate that despite retaining basal respiration, OsiR cells display clear hallmarks of mitochondrial dysfunction, including reduced ATP production, increased proton leak, and diminished respiratory flexibility. These changes functionally validate the molecular rewiring of mitochondrial programs observed in OsiR cells and position mitochondrial remodeling as a core feature of acquired resistance to osimertinib.

### 2.5 Exometabolomic Profiling reveals biomarkers of Osimertinib resistance

To further characterize the metabolic reprogramming of OsiR cells, we performed exometabolomic profiling of culture supernatants to identify the metabolites released into the extracellular medium, using a combination of HPLC-DAD-FD, GC-MS, and LC-HRMS approaches. This analysis enabled the identification of metabolites secreted into the extracellular environment and provided insights into pathway-level alterations. Pathway and biomarker enrichment analyses revealed significant dysregulation of central carbon metabolism in OsiR cells, consistent with transcriptomic and bioenergetic findings. Specifically, OsiR cells exhibited altered flux through glycolysis, gluconeogenesis, and the pentose phosphate pathway (PPP). Glyceraldehyde-3-phosphate (G3P) levels were reduced, while 3-phosphoglyceric acid (3-PGA), pyruvate, lactate, acetaldehyde, and acetate were markedly elevated (**Figure 5a**). Notably, acetate, undetectable in Par cells, emerged as the predominant acidic metabolite in OsiR cells and, together with lactate, contributed to ECM acidification. These findings align with the enhanced glycolytic phenotype observed in transcriptomic datasets and were functionally corroborated by rapid acidification of the cell culture medium (**Figure 5b**), a metabolic hallmark of aggressive cancer cells.

**Figure 5.**
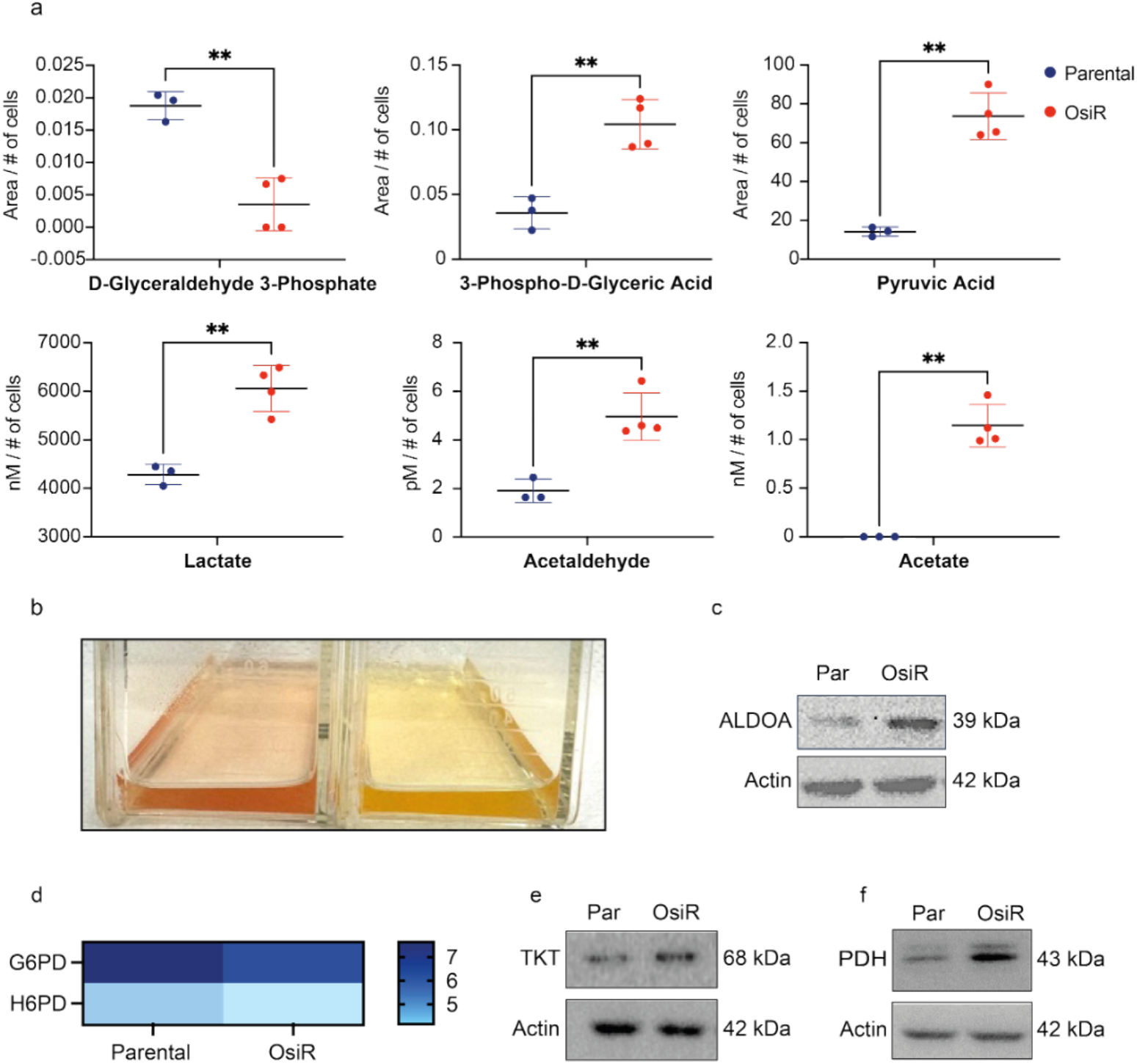
Exometabolomic profiling uncovers biomarkers of Osimertinib-resistance in EGFR mutant NSCLC cells. **(a)** Exometabolomic analysis of secreted metabolites in the cell culture medium of H1975 Par (blue) and OsiR cells (red). Error bars represent standard deviation (SD); p-values are indicated (**p<0.002). **(b)** Representative images of the extracellular medium from H1975 OsiR (right) and Par cells (left). Yellow coloration indicates increased acidification in OsiR cells. **(c)** Western blot analyses of H1975 Par and OsiR cells. Protein lysates were immunoblotted with an anti-ALDOA antibody. Loading was assessed with an anti-β-actin antibody. Expected size is shown in kDa. **(d)** Heat map showing relative gene expression of G6PD and H6PD in Par and OsiR H1975 cells. **(e-f)** Western blot analyses of Par and OsiR cells. Protein lysates were immunoblotted with an anti-TKT or PDH antibody.

### Linking the Warburg Effect and Alternative NADPH Production

The accumulation of pyruvate and lactate in OsiR cells points to a shift toward aerobic glycolysis, a hallmark of the Warburg effect, where cancer cells preferentially rely on glycolysis over OXPHOS, even in the presence of oxygen, to rapidly generate ATP and biosynthetic precursors.

Western blot analysis confirmed upregulation of the glycolytic enzyme Aldolase A (ALDOA) in OsiR cells (**Figure 5c**). ALDOA catalyzes the cleavage of fructose-1,6-bisphosphate to glyceraldehyde-3-phosphate and dihydroxyacetone phosphate (DHAP), enhancing glycolytic flux while supplying key intermediates for nucleotide, amino acid, and lipid biosynthesis. In this context, G3P and DHAP function not merely as intermediates en route to pyruvate but as key biosynthetic nodes. Instead of proceeding through glycolysis to form pyruvate and fuel the mitochondria for energy production via the TCA cycle, G3P in OsiR cells appears redirected into the non-ox-PPP, bypassing the oxidative arm that generates NADPH.

Transcriptomic analysis revealed significant downregulation of the ox-PPP enzymes, including glucose-6-phosphate dehydrogenase (G6PD) (−0.57; adj. p-value 0.0003) and hexose-6-phosphate dehydrogenase (H6PD) (−0.81; adj. p-value 9.34E-08) (**Figure 5d**). In contrast, the non-ox PPP enzyme transketolase (TKT) was upregulated (**Figure 5e**), indicative of a metabolic bias toward a biosynthetic PPP program that functions independently of the ox-PPP, and instead supplies structural intermediates rather than redox equivalents.

Given the suppression of the oxidative PPP, we hypothesized that OsiR cells activate a novel alternative route for cytosolic NADPH generation via the **pyruvate-acetaldehyde-acetate (PAA) pathway**. In this pathway, pyruvate is sequentially converted into acetaldehyde and acetate, with NADPH generated during acetaldehyde detoxification. This bypass enables redox balancing while preserving glycolysis as the main ATP source, consistent with a Warburg-like metabolic state.

This rerouting ensures that glucose-derived carbon is funneled exclusively through glycolysis, avoiding diversion into the ox-PPP. In parallel, the PAA pathway allows pyruvate to generate both NADPH for redox homeostasis and cytoplasmic acetyl-CoA for fatty acid and membrane biosynthesis. Detoxification of acetaldehyde into acetate is mediated by NADP^+^-dependent aldehyde dehydrogenases, ALDH2 (mitochondrial) and ALDH7A1 (cytoplasmic), both of which were transcriptionally upregulated in OsiR cells. Western blot validation confirmed increased expression of ALDOA (**Figure 5c**), mitochondrial pyruvate dehydrogenase (PDH) (**Figure 5f)**, succinate dehydrogenase (SDH) **(Figure 3c)**, TKT (**Figure 5e**), and ALDH2/ALDH7A1 (**Figure 1c**), supporting a coordinated metabolic adaptation that favors biosynthetic output and redox adaptability in drug-resistant cells.

## DISCUSSION

Resistance to EGFR-targeted tyrosine kinase inhibitors (TKIs) remains a major challenge in treating NSCLC. While osimertinib is currently the standard of care for EGFR-mutant NSCLC, its clinical efficacy is ultimately limited by the emergence of acquired resistance. Although mechanisms such as secondary mutations, lineage plasticity, and epigenetic reprogramming are well-documented, the contribution of mitochondrial adaptation to resistance remains largely uncharted.

Here, we uncover a mitochondrial-centric dimension of osimertinib resistance, characterized by both structural and metabolic reprogramming. Resistant cells show fragmented mitochondrial networks, impaired OXPHOS, and a collapse of spare respiratory capacity. These features coincide with reduced activity in the oxidative pentose phosphate pathway, and a shift toward aerobic glycolysis, a phenotype consistent with the Warburg effect. These alterations converge into a dysfunctional mitochondrial state that paradoxically supports anabolic activity, redox homeostasis, and survival.

We introduce the term *mitochondriomics* to capture this complex and systems-level mitochondrial reprogramming, encompassing genomic, transcriptomic, structural, and metabolic changes that enable therapeutic escape. By integrating transcriptomic, exometabolomic, real-time carbon flux tracing, high-resolution imaging and structural analyses, we demonstrate that mitochondria serve as both sensors of drug-induced stress and effectors of adaptive responses.

Mechanistically, OsiR cells harbor mutations in both mitochondrial and nuclear-encoded ETC components, including assembly factors. Some mutations pre-exist at low allelic frequency, while others emerge *de novo*, supporting a dual model of clonal selection and drug-induced evolution. The convergence of mutations on key ETC components suggests functional selection rather than random genetic drift. Functionally, the constellation of mutations observed in resistant cells is accompanied by reduced coupling efficiency, increased proton leak and diminished respiratory plasticity.

Structurally, resistant cells accumulate donut-shaped and spherical mitochondria, morphologies linked to oxidative stress and impaired fission-fusion dynamics(24) (25). These changes are tightly coupled to a glycolytic shift*(26)* and increased dependence on cytosolic metabolic pathways for ATP, NADPH, and biosynthetic intermediates.

Biochemically, mitochondria in resistant cells undergo major metabolic detours. Most notably, we identify a non-canonical, functionally repurposed route: the pyruvate-acetaldehyde-acetate (PAA) pathway, which displays remarkable flexibility and functions as a *metabolic toolkit* for endurance. This insight emerged from a retrospective, data-driven analysis that combined computational mining with manual curation of metabolic gene expression. This approach highlights the limitations of traditional pathway-centric models, which overlooked this adaptive, non-canonical route, and underscored the value of systems integration.

Based on the upregulation of PDHA1, which encodes the E1 alpha subunit of the pyruvate dehydrogenase (PDH) complex, and of mitochondrial ALDH2 and cytosolic ALDH7A1, enzymes responsible for oxidizing acetaldehyde to acetate, we propose a functional metabolic shunt. Specifically, we suggest that pyruvate is rerouted through an atypical decarboxylation step, enabled by neomorphic PDH activity, reminiscent of yeast pyruvate decarboxylase, to form acetaldehyde. Acetaldehyde is then oxidized by ALDH2 and ALDH7A1 into acetate. This rerouted pathway forms what we term the PAA bypass.

This shift serves a dual adaptive purpose: detoxifying acetaldehyde and regenerating cytosolic NADPH, which is essential for maintaining redox homeostasis and supporting anabolic metabolism under mitochondrial stress. Notably, these compensatory functions emerge in a context where canonical NADPH-generating pathways, such as the oxPPP, are suppressed. Despite the marked downregulation of G6PD and H6PD, key oxPPP enzymes, resistant cells maintain NADPH and acetyl-CoA production via the PAA bypass. Thus, OsiR cells adopt this alternative, ALDH-dependent NADPH production system, repurposing components of the PAA pathway to sustain redox balance and biosynthetic capacity when OXPHOS is compromised.

The PAA shunt, while distinct, shares similarities with the Warburg effect and is characterized by enhanced glycolytic flux and the accumulation of pyruvate, lactate, acetaldehyde, and acetate. While previous studies have reported altered pyruvate metabolism (27), acetate formation from acetaldehyde (28), and TCA reshaping in cancer (29), our data uniquely position the PAA pathway as a defined metabolic strategy to sustain NADPH and acetyl-CoA pools, under mitochondrial constraint.

Notably, ALDH upregulation has previously been linked to cancer stemness, therapeutic resistance, and poor prognosis (30). Here, its role expands to include cytosolic NADPH generation and metabolic resilience. The upregulation of antioxidant defenses, including glutathione peroxidases and superoxide dismutases, reinforces the importance of redox buffering in sustaining the resistant phenotype.

In parallel, increased expression of ALDOA and TKT suggests compensatory activation of the non-oxidative branch of the PPP. This adaptation supports the increased glycolytic flux in resistant cells and redirects carbon flow into the non-oxPPP to sustain anabolic needs. ALDOA, a key glycolytic enzyme, accelerates the conversion of fructose-1,6-bisphosphate into triose phosphates, which serve as substrates for the non-oxidative PPP. TKT, which is also upregulated, facilitates the interconversion of sugar phosphates, thereby fueling the biosynthesis of nucleotides and amino acids. TKT plays a critical role in supplying ribose-5-phosphate and other anabolic precursors.

Overall, this metabolic network reinforces a functional division of labor and highlights the complementarity between the PAA axis and non-oxPPP: while the PAA bypass restores cytosolic NADPH and acetyl-CoA under mitochondrial stress, the non-oxPPP sustains biosynthetic outputs necessary for proliferation, even in the context of impaired mitochondrial function.

Together, our data redefine osimertinib resistance as a multidimensional program involving structural, genetic, and metabolic mitochondrial remodeling. Mitochondrial dysfunction emerges as an adaptive engine for survival. ETC mutations, increased proton leak, reduced spare respiratory capacity, and activation of the PAA bypass form an integrated strategy for endurance that allows resistant cells to maintain energy production, redox homeostasis, and biomass synthesis.

Therapeutically, this expanded view of resistance suggests new mitochondria-centric vulnerabilities, advocating that targeting components of this adaptive network may re-sensitize resistant cells. Combinatorial strategies using OXPHOS inhibitors (e.g., metformin) alongside TKIs have shown promise in delaying resistance onset in preclinical studies (31).

In conclusion, our work introduces a new conceptual framework for understanding resistance through the lens of **mitochondriomics**. By decoding mitochondrial adaptive transformation, metabolic rewiring, fission-fusion dynamics, and stress adaptation, we position mitochondria as a central hub in therapeutic escape. This integrative perspective offers an integrated paradigm to decode resistance mechanisms and informs the development of novel mitochondria-targeted therapeutic strategies. Importantly, mitochondrial dysfunction may serve as a predictive biomarker of resistance, guiding personalized approaches for patients with EGFR-mutant NSCLC.

## MATERIAL AND METHODS

### Cell culture conditions

The parental human lung cancer cell line H1975, which harbors the EGFR L858R/T790M mutation, was obtained from ATCC. The osimertinib-resistant variant was generated by gradually exposing H1975 parental cells to increasing concentrations of osimertinib, up to a final concentration of 1 μM, at which the cells remained viable. Cells were maintained in RPMI-1640 medium supplemented with 10% fetal bovine serum (FBS), 1% L-glutamine, and 1% penicillin/streptomycin, at 37°C in a humidified incubator with 5% CO2.

### Total RNA Sequencing

Total RNA was extracted from both H1975-Par and -OsiR cells using the RNeasy Plus Mini Kit (QIAGEN#74134), following the manufacturer’s instructions. RNA concentration and integrity were evaluated using a NanoDrop 2000 spectrophotometer (Thermo Fisher Scientific). RNA sequencing was performed by IGATech (IGA Technology Services, Udine, Italy). RNA libraries were prepared using the Universal Plus mRNA-Seq kit (Tecan Genomics, Redwood City, CA), following the manufacturer’s protocol for library type *fr-secondstrand*. RNA quality was evaluated using either the Agilent 2100 Bioanalyzer RNA assay (Agilent Technologies, Santa Clara, CA) or Caliper LabChip GX system (PerkinElmer, Waltham, MA). Final libraries were quantified with a Qubit 2.0 Fluorometer (Invitrogen, Carlsbad, CA) and their quality was verified using the Agilent Bioanalyzer DNA assay or Caliper. Sequencing was performed using a NovaSeq 6000 platform (Illumina, San Diego, CA) with paired-end 150 bp reads. The number of reads (in millions) generated per sample is provided below:

P1: 120.08

P2: 116.47

P3: 114.88

OR1: 120.35

OR2: 115.34

OR3: 103.65

Base calling, demultiplexing, and adapter masking were performed using Illumina BCL Convert v3.9.31. Adapter sequences were masked during demultiplexing, with masked bases converted to “N” characters and associated quality scores overwritten to a value of 2 to facilitate compatibility with standard trimming and quality filtering tools.

### RNA sequencing reads alignment and count

RNA sequencing reads were aligned to the genome assembly GRCh38p14 using the aligner STAR(32), resulting in a mean of uniquely mapped reads per sample of 92.4% (SD 2.5%). Mapped reads were counted using featureCounts(33) and annotation file “Homo_sapiens.GRCh38.110.gtf” from Ensembl (https://www.ensembl.org), resulting in a mean of assigned reads per sample of 90.8% (SD 0.2%). The genes-by-samples count matrix was filtered to keep only genes with at least 3 counts per million (cpm) in 3 samples, a constraint fulfilled by 11.298 genes.

### Differential Gene Expression Analysis

Samples were normalized using the TMM (trimmed mean of M values) method(34) available in the R package edgeR(35). Differential gene expression (DGE) analysis between H1975-Par and H1975-OsiR cells was performed using limma(36). Genes with an absolute log fold change (|LogFC|) ≥ 0.25 and a false discovery rate (FDR) ≤ 0.05 were considered significantly differentially expressed.

### Enrichment Analysis

The list of significantly upregulated genes was analyzed using the EnrichR platform for functional enrichment (37) (38, 39). Enrichment analysis was performed across multiple databases, including Gene Ontology (GO) categories, Biological Processes (BP) and Cellular Components (CC), as well as Hallmark (HM) and WikiPathway (WP), to identify overrepresented biological pathways and cellular functions.

### Cell Collection and Protein Extraction

H1975 Par and OsiR cells were detached from culture plates using trypsin-EDTA, centrifuged at 261x g for 5 minutes at room temperature, and washed with PBS. The resulting cell pellet was snap-frozen in liquid nitrogen and stored at −80°C until further processing. To extract proteins, the cell pellet was lysed for 30 minutes in 1X Triton X-100 (Thermo Scientific, #A16046) supplemented with cOmplete EDTA-free Protease Inhibitor Cocktail and PhosStop Phosphatase Inhibitor Cocktail. The lysates were incubated on ice for 30 minutes with vortexing every 10 minutes. After incubation, the samples were centrifuged at 12,000 × g for 20 minutes at 4°C, and the supernatant was collected and stored at - 80°C.

### SDS-PAGE and Western Blotting

For protein analysis, 15 μg of protein from each sample (Par and OsiR cells) was separated by SDS-PAGE on 10% gel and subsequently transferred to nitrocellulose membranes with the TransBlot Transfer System (Bio-Rad). Membranes were blocked for 1 hour at room temperature in either 5% non-fat dry milk or 5% bovine serum albumin (BSA) diluted in Tris-buffered saline with 0.1% Tween-20 (1X TBS-T). Primary antibodies (listed below) were incubated overnight at 4°C. After incubation, membranes were washed three times for 10 minutes each with 1X TBS-T at room temperature. HRP-conjugated secondary antibodies (listed below) were then applied for 1 hour at room temperature. Proteins bands were detected by chemiluminescence (Pierce ECL Western Blotting Substrate Cat #32106) and visualized using the Chemidoc Imaging System (Biorad). Membranes were subsequently stripped using Stripping Buffer Solution (Himedia, #ML163), as per the manufacturer’s instructions, and re-probed with an anti-β-actin mouse antibody to verify equal protein loading. After three washes with 1X TBS-T, membranes were incubated with the appropriate anti-mouse secondary antibody (details below).

| ANTIBODIES AND DILUTIONS |  |
| --- | --- |
| Primary Antibody | Dilution |
| anti-β-actin mouse antibody (Santa Cruz #C3022) | 1/1000 (in 5% non fat milk) |
| anti-ALDOA rabbit antibody (Antibodies #A97723) | 1/1000 (in 5% non fat milk) |
| anti-ALDH2 rabbit antibody (Antibodies #A13393) | 1/1000 (in 5% non fat milk) |
| anti-TKT rabbit antibody (Antibodies #A15167) | 1/1000 (in 5% non fat milk) |
| anti-ALDH7A1 rabbit antibody (Invitrogen #PA5-29974) | 1/1000 (in 5% non fat milk) |
| anti-MFN2 rabbit antibody (Cell Signaling Technology #9482) | 1/1000 (in 5% non fat milk) |
| anti-DRP1 rabbit antibody (Cell Signaling Technology #8570) | 1/1000 (in 5% BSA) |
| anti-COX IV rabbit antibody (Cell Signaling Technology #4850) | 1/1000 (in 5% BSA) |
| anti-CytC rabbit antibody (Cell Signaling Technology #4280) | 1/1000 (in 5% non fat milk) |
| anti-PHB1 rabbit antibody (Cell Signaling Technology #2426) | 1/1000 (in 5% BSA) |
| anti-PDHA1 rabbit antibody (Cell Signaling Technology #3205) | 1/1000 (in 5% BSA) |
| anti-SDHA rabbit antibody (Cell Signaling Technology #11998) | 1/1000 (in 5% BSA) |
| anti-VDAC rabbit (Cell Signaling Technology #4661) | 1/1000 (in 5% BSA) |
| Secondary Antibody | Dilution |
| anti-mouse (Immunoreagents #NC27606) | 1/2000 (in 5% non fat milk) |
| anti-rabbit (Cell Signaling Technologies #7074) | 1/2000 (in 5% non fat milk) |

### Mitochondrial Staining and Imaging

Mitochondrial staining was performed using the MITO-ID Red Detection Kit (Enzo Life Sciences, Cat # ENZ-51007) according to the manufacturer’s protocol. Confocal imaging of stained cell samples was carried out using a Zeiss LSM 900 Airyscan2 (Carl Zeiss, Jena, Germany). A 63x Apochromat oil-immersion objective (NA = 1.4) was employed, and the pinhole aperture of the confocal system was set at 1 airy unit. The excitation wavelength was set to 561 nm, and emission was detected in the 565-625 nm range. Image stacks were acquired at a resolution of 1596 × 1596 pixels^2^, corresponding to a field of view of 78.0 × 78.0 µm^2^, with an axial step of 0.17 µm. Four averages were taken per line scan to ensure signal clarity. The Airyscan system provides an optical lateral resolution of approximately 120 nm and an axial resolution of about 350 nm, enabling precise visualization of mitochondrial morphology.

### Image Analysis

3D mitochondrial reconstruction was performed by Airyscan confocal microscopy.

Image processing was performed using the *Mitochondria Analyzer* plugin(40) integrated into the open-source software Fiji (NIH, Bethesda, USA). The pipeline enables 3D analysis, allowing for detailed quantification of mitochondrial morphology across multiple optical planes. The following morphological parameters were assessed: Mitochondrial number per cell, Mean mitochondrial volume, Mean mitochondrial surface area, Mitochondrial sphericity, Number of branches per mitochondrion, Total branch length per mitochondrion, Branch junctions per mitochondrion and Branch end-points per mitochondrion.

### Mitochondria Isolation and Extraction of Mitochondrial DNA (mtDNA) for Sequencing

Mitochondria were isolated from both H1975-Par and H1975-OsiR cells using the Qproteome Mitochondrial Isolation kit (QIAGEN#37612), according to the manufacturer’s instructions. Mitochondrial DNA (mtDNA) was then extracted from the isolated mitochondria employing the DNA Micro kit (QIAGEN#56304) by following manufacturer’s instructions. mtDNA concentration and purity were assessed using the DS-11 spectrophotometer (DeNovix), the Quant-iT™ dsDNA High Sensitivity Assay Kit (Thermo Fisher Scientific), and capillary electrophoresis (Fragment Analyzer, Agilent) to ensure DNA integrity. A total of 300 ng of high-quality DNA was sheared to a target size range of 7–12 kb and used for library preparation with the SMRTbell Express Template Prep Kit v3.0 (Pacific Biosciences), according to the manufacturer’s instructions. Library preparation included DNA damage repair, end repair with A-tailing, and ligation of barcoded SMRTbell adapters. The resulting libraries were purified using SMRTbell cleanup beads (PacBio), and size and concentration were re-evaluated using capillary electrophoresis and fluorometric assays. SMRTbell libraries were pooled by equal mass and submitted to high-throughput sequencing at the Genesupport SA facility (Plan-les-Ouates, CH). Sequencing was performed by Fasteris (Genesupport SA Life Sciences, Switzerland) on the PacBio Sequel IIe platform, using the Sequel Sequencing Kit 3.0 with 1800-minute movies, and SMRT Link Software v11.0 for HiFi read generation, demultiplexing, and primary analysis.

### Mitochondrial DNA sequencing, variant call and annotation

mtDNA sequencing was performed using the PacBio HiFi platform to ensure high-fidelity detection of single nucleotide variants and structural alterations. Long-reads from Par and OsiR samples were aligned to the Revised Cambridge Reference Sequence (rCRS) of the Human Mitochondrial DNA (GenBank sequence “NC_012920”) using pbmm2 (https://github.com/PacificBiosciences/pbmm2), a PacBio-optimized frontend for the *minimap2* aligner(41). Variant calling was performed using MutServe2(42). The analysis detected a total of 2662 SNVs in Par cells and 2416 SNVs in OsiR cells. Variants were annotated using Ensembl Variant Effect Predictor (VEP, https://www.ensembl.org/Tools/VEP). Only variants with AF ≥ 0.01 (in Par or OsiR cells), AF fold change (OsiR/Par and Par/OsiR) ≥ 1.5, and classified by VEP IMPACT as MODIFIER or MODERATE or HIGH were kept. This analysis led to a total of 171 variants in genes that encode mitochondrial respiratory chain complexes (annotation from MitoCarta3.0(43) **(Supplementary Data 1)**.

### Genomic DNA extraction and whole genome sequencing

Total DNA was extracted from H1975-Par and H1975-OsiR cells using the DNA Micro Kit (QIAGEN, #56304), according to the manufacturer’s instructions. DNA concentration and purity were assessed by spectrophotometry using a NanoDrop 2000 (Thermo Fisher Scientific). Genomic DNA was used for whole genome sequencing (WGS), which was performed by IGATech (IGA Technology Services, Italy). Libraries were prepared using the Celero™ DNA-Seq kit (NuGEN, San Carlos, CA) for high-quality genomic DNA, following the manufacturer’s instructions. Final libraries were quantified by Qubit 2.0 Fluorometer (Invitrogen, Carlsbad, CA) and quality-checked with the Agilent 2100 Bioanalyzer High Sensitivity DNA assay (Agilent Technologies, Santa Clara, CA) to assess fragment size distribution and concentration, ensuring suitability for sequencing. Sequencing was performed on a NovaSeq 6000 platform (Illumina) in paired-end 150 mode, generating 150 bp paired-end reads. Approximately 90 Gbp of data were obtained per sample, corresponding to ∼30X coverage of the human genome. Base calling, demultiplexing, and adapter masking were performed using BCL Convert v3.9.3.

### Whole genome sequencing reads alignment, variant call and annotation

DNA sequencing reads alignment was performed with BWA-MEM (https://bio-bwa.sourceforge.net) against the human assembly GRCh38p14. Aligned reads were analyzed with GATK (cleaning, reordering, sorting, fixing mate information, marking duplicates and base recalibration) (44). Mutations in OsiR vs Par, and in Par vs OsiR, were called using GATK Mutect2 and filtered using GATK “FilterMutectCalls”. Variants were annotated using Ensembl Variant Effect Predictor (VEP, https://www.ensembl.org/Tools/VEP). Only variants classified by VEP IMPACT as MODIFIER or MODERATE or HIGH were kept. This analysis led to a total of 121 variants in nuclear genes that encode mitochondrial respiratory chain complexes (annotation from MitoCarta3.0(43)) **(Suppl. Data 2)**.

### Seahorse Metabolic Flux Analysis

Real-time metabolic flux analysis was performed using the XFe96 Extracellular Flux Analyzer (Seahorse Bioscience). H1975P and OsiR cells were seeded in Seahorse XF96 cell culture plate at a density of 15,000 cells per well. Prior to the assay, cells were washed and incubated in Seahorse XF Base Medium supplemented with 10 mM glucose, 1 mM pyruvate, and 2 mM L-glutamine. Plates were maintained in a non-CO_2_ incubator at 37°C for 1 hour prior to allow temperature and pH equilibration. Oxygen consumption rates (OCRs) were measured under basal conditions and following the sequential injection of mitochondrial inhibitors, following the manufacturer’s instructions. Final concentrations of the inhibitors were as follows: 1.5µM oligomycin, 1µM carbonyl cyanide-4-phenylhydrazone (FCCP), 0.5µM antimycin A, and 0.5µM rotenone.

#### Cell culture medium collection for exometabolome profilin

H1975P and OsiR cells were cultured in at least three biological replicates per condition for 72 hours at 37°C in a humidified incubator with 5% CO_2_, until reaching 90-100% confluence. The cell culture medium, containing the exometabolome, was then collected for further processing. Cells were detached and counted using a hemocytometer to enable normalization of metabolite levels.

#### Pyruvate Quantification by UHPLC-HRMS

Pyruvate levels were quantified by LC-HRMS. Analyses were carried out on an ultra-High-Performance Liquid Chromatography system (UHPLC; Vanquish Flex Binary pump) coupled with a diode array detector (DAD) and a high resolution (HR) Q Exactive Plus MS, based on Orbitrap technology, equipped with a heated electrospray ionization (HESI) source (Thermo Fischer Scientific Inc., Bremem, Germany).

Chromatographic separation was achieved on a Zorbax® Eclipse Plus Phenyl-Hexyl Column (4.6 × 250 mm, 5 µm; Agilent Technologies, CA, USA) with a flow rate of 0.8 mL/min and a 1:1 split between the HRMS and DAD/UV detectors. The injections volume was 2 μL, and the column temperature was maintained at 40°C. Eluents consisted of MeOH/HCOOH 0.5% v/v (solvent B) and H_2_O/HCOOH 0.5% v/v (solvent A), both of ultra-purity grade (ROMIL Ltd, Cambridge, UK). The gradient was as follows: 0-15 min: isocratic 100% A; 15-25 min: linear gradient from 0 to 80% B; 25-35 min: linear gradient from 80% to 100% B; 35-37 min (2 minutes): isocratic 100% B.; 37-44 min: re-equilibration with 100% solvent A.

#### HRMS Parameters

Ionization was performed in both positive and negative modes: nebulization voltage 3400 V (+) and 3200 V(−), capillary temperature 290°C, sheath gas (N_2_) 24 (+) and 28 (−) arbitrary units, auxiliary gas (N_2_) 5 (+) and 4 (−) arbitrary units, S lens RF level 50. Full scan mode was used over an m/z range of 54-800, with a resolution of 70,000 at m/z 200. Data acquisition and analysis were conducted using Xcalibur 3.1 software (Thermo Fischer Scientific Inc., Bremen, Germany). LCMS files were processed using Xcalibur Version 4.1.0.0. Extracted Ion Chromatograms (EIC) were obtained by exact mass ranges (calculated from molecular formula C_n_H_n_O_n_P_a_N_n_S_n_, -H for negative ions, +H for positive ions) with ± 3 ppm (ΔM/M) *10^6^ accuracy. Chromatogram peaks were manually integrated, and quantitative analysis was carried out in Microsoft Excel.

#### Lactate Quantification by RP-HPLC-DAD

Due to its high extracellular concentration (mM) saturating HRMS detectors, lactate was quantified by reversed-phase HPLC (RP-HPLC) with diode array detection (HPLC-DAD)^57^. For RP-HPLC-DAD analysis, samples were diluted 1:5 times in 5 mM sulfuric acid, filtered using 0.20 μm RC Mini-Uniprep filters (Agilent Technologies, Italy), and injected (V_inj_= 5μL) into the same column employed for LC-HRMS analysis. Metabolite identification was based on comparison with standard retention times and UV spectra. Quantification was performed via calibration curves generated from analytical standards, and peaks were manually integrated.

#### Acetate and Acetaldehyde Quantification by Headspace GC-MS

Acetate and acetaldehyde were quantified by head space gas chromatography-mass spectrometry (HS-GC-MS). Acetaldehyde was measured directly, while acetate required derivatization with trialkyloxonium salts, since LCMS is not accurate in the determination of low MW metabolites. For acetate analysis, 500 μL of sample or standard solution (0, 50, 100, 250, 400 μg/mL from Certified Standard), 50 μL of ^2^H_3_ acetate at 1000 μg/mL (Internal Standard), and 100 μL of 5 M H_2_SO_4_ were added to 10 mL HS vials and sealed immediately. Standards and samples were prepared in triplicate. Calibration curves for quantification were prepared by plotting the areas at m/z 43 /area at m/z 46 ratio vs [acetate]/[acetate ^2^H_3_] concentration ratio, using Agilent MassHunter Quantitative Analysis (v10.2). Measurements were performed using an Agilent 6850 GC system equipped with a 5975c single quadrupole MS detector and a GC 80 CTC PAL ALS autosampler. Separations were performed in J&W DB-WAX-UI capillary columns (30.0 m, 250.00 μm internal diameter, 0.50 μm film thickness) at constant flow of 1.0 mL/min (average velocity: 36 cm/s; incubation temperature 80 °C; incubation time 600 s; 1000 μL injection volume). The oven temperature program was: initial temperature 50 °C (hold 5 min), ramp up to 150°C at 10°C/min (hold 2 min); ramp to 240°C at 20°C/min (hold 8.5 min); total run time 30 min. Pulsed Splitless inlet mode; initial temperature 200 °C, pressure 45.4 kPa, pulse pressure 120 kPa, pulse time 0.10 min, purge flow 200.0 mL/min, purge time 0.10 min, total flow 203.7 mL/min, gas type helium; transfer line temperature 250 °C; SIM acquistion mode: EI ionization mode at 70eV electron energy; Ion Source Temp. 250°C; Quadrupole Temperature: 150°C; SIM mode was used with acquisition at m/z 43, 46, 60, and 63 (Dwell Time 100 ms). Quantifier ions: 43 and 46; qualifier ions: 60 and 63. Data acquisition was managed via MSD ChemStation (Agilent, vE.02.02).

## Data Availability

RNA sequencing, mitochondrial DNA sequencing and Whole genome sequencing data were deposited into EMBL-EBI’s ArrayExpress under accession number,,, respectively.

## FUNDING

This work was supported by the AIRC Investigator Grant 2021 (ID 25734), PNNR THE Spoke 1 Award, the PNRR-MCNT1-2023-12377671 Grant, the ELMO Pisa Foundation Grant, the FPS Grant 2024, and private donations from the Gheraldeschi and Pecoraro families to EL. AC was supported by the European Union - Next Generation EU, Mission 4 Component 2, project “Strengthening BBMRL.it”. AA acknowledges funding from A-0001263-00-00.

## Notes

### Competing Interest Statement

The authors have declared no competing interest.

## REFERENCES

1. Levantini E, Maroni G, Del Re M, Tenen DG. EGFR signaling pathway as therapeutic target in human cancers. Semin Cancer Biol. 2022. Epub 2022/04/16. doi: 10.1016/j.semcancer.2022.04.002. PubMed PMID: 35427766.

2. Ali A, Levantini E, Teo JT, Goggi J, Clohessy JG, Wu CS, Chen L, Yang H, Krishnan I, Kocher O, Zhang J, Soo RA, Bhakoo K, Chin TM, Tenen DG. Fatty acid synthase mediates EGFR palmitoylation in EGFR mutated non-small cell lung cancer. EMBO Mol Med. 2018;10(3). Epub 2018/02/17. doi: 10.15252/emmm.201708313. PubMed PMID: 29449326; PMCID: PMC5840543.

3. Ali A, Levantini E, Fhu CW, Teo JT, Clohessy JG, Goggi JL, Wu CS, Chen L, Chin TM, Tenen DG. CAV1 - GLUT3 signaling is important for cellular energy and can be targeted by Atorvastatin in Non-Small Cell Lung Cancer. Theranostics. 2019;9(21):6157–74. Epub 2019/09/20. doi: 10.7150/thno.35805. PubMed PMID: 31534543; PMCID: PMC6735519.

4. Warburg O. On the origin of cancer cells. Science. 1956;123(3191):309–14. Epub 1956/02/24. doi: 10.1126/science.123.3191.309. PubMed PMID: 13298683.

5. Diaz-Ruiz R, Rigoulet M, Devin A. The Warburg and Crabtree effects: On the origin of cancer cell energy metabolism and of yeast glucose repression. Biochimica et Biophysica Acta (BBA) - Bioenergetics. 2011;1807(6):568–76. doi: 10.1016/j.bbabio.2010.08.010.

6. Ferreira LM. Cancer metabolism: the Warburg effect today. Exp Mol Pathol. 2010;89(3):372–80. Epub 2010/09/02. doi: 10.1016/j.yexmp.2010.08.006. PubMed PMID: 20804748.

7. Medes G, Paden G, Weinhouse S. Metabolism of neoplastic tissues. XI. Absorption and oxidation of dietary fatty acids by implanted tumors. Cancer Res. 1957;17(2):127–33. Epub 1957/02/01. PubMed PMID: 13413849.

8. Schug ZT, Vande Voorde J, Gottlieb E. The metabolic fate of acetate in cancer. Nat Rev Cancer. 2016;16(11):708–17. Epub 2016/10/25. doi: 10.1038/nrc.2016.87. PubMed PMID: 27562461; PMCID: PMC8992383.

9. Robey RB, Hay N. Mitochondrial hexokinases, novel mediators of the antiapoptotic effects of growth factors and Akt. Oncogene. 2006;25(34):4683–96. Epub 2006/08/08. doi: 10.1038/sj.onc.1209595. PubMed PMID: 16892082.

10. Liberti MV, Locasale JW. The Warburg Effect: How Does it Benefit Cancer Cells? Trends in Biochemical Sciences. 2016;41(3):211–8. doi: 10.1016/j.tibs.2015.12.001.

11. Colombaioni L, Onor M, Benedetti E, Bramanti E. Thallium stimulates ethanol production in immortalized hippocampal neurons. PLoS One. 2017;12(11):e0188351. Epub 2017/11/22. doi: 10.1371/journal.pone.0188351. PubMed PMID: 29161327; PMCID: PMC5697870.

12. Bramanti E, Onor M, Colombaioni L. Neurotoxicity Induced by Low Thallium Doses in Living Hippocampal Neurons: Evidence of Early Onset Mitochondrial Dysfunction and Correlation with Ethanol Production. ACS Chem Neurosci. 2019;10(1):451–9. Epub 2018/10/23. doi: 10.1021/acschemneuro.8b00343. PubMed PMID: 30346713.

13. Campanella B, Colombaioni L, Nieri R, Benedetti E, Onor M, Bramanti E. Unraveling the Extracellular Metabolism of Immortalized Hippocampal Neurons Under Normal Growth Conditions. Front Chem. 2021;9:621548. Epub 2021/05/04. doi: 10.3389/fchem.2021.621548. PubMed PMID: 33937186; PMCID: PMC8085660.

14. Colombaioni L, Campanella B, Nieri R, Onor M, Benedetti E, Bramanti E. Time-dependent influence of high glucose environment on the metabolism of neuronal immortalized cells. Anal Biochem. 2022;645:114607. Epub 2022/03/02. doi: 10.1016/j.ab.2022.114607. PubMed PMID: 35227660.

15. Wu J, Chen Y, Lin Y, Lan F, Cui Z. Cancer-testis antigen lactate dehydrogenase C4 as a novel biomarker of male infertility and cancer. Front Oncol. 2022;12:936767. Epub 2022/11/22. doi: 10.3389/fonc.2022.936767. PubMed PMID: 36408133; PMCID: PMC9667869.

16. Peng W, Chen J, Xiao Y, Su G, Chen Y, Cui Z. Cancer-Testis Antigen LDH-C4 in Tissue, Serum, and Serum-Derived Exosomes Serves as a Promising Biomarker in Lung Adenocarcinoma. Front Oncol. 2022;12:912624. Epub 2022/07/12. doi: 10.3389/fonc.2022.912624. PubMed PMID: 35814471; PMCID: PMC9263124.

17. Muzio G, Maggiora M, Paiuzzi E, Oraldi M, Canuto RA. Aldehyde dehydrogenases and cell proliferation. Free Radic Biol Med. 2012;52(4):735–46. Epub 2011/12/31. doi: 10.1016/j.freeradbiomed.2011.11.033. PubMed PMID: 22206977.

18. Dinavahi SS, Bazewicz CG, Gowda R, Robertson GP. Aldehyde Dehydrogenase Inhibitors for Cancer Therapeutics. Trends Pharmacol Sci. 2019;40(10):774–89. Epub 2019/09/14. doi: 10.1016/j.tips.2019.08.002. PubMed PMID: 31515079.

19. Schagger H, Link TA, Engel WD, von Jagow G. Isolation of the eleven protein subunits of the bc1 complex from beef heart. Methods Enzymol. 1986;126:224–37. Epub 1986/01/01. doi: 10.1016/s0076-6879(86)26024-3. PubMed PMID: 2856130.

20. Timon-Gomez A, Nyvltova E, Abriata LA, Vila AJ, Hosler J, Barrientos A. Mitochondrial cytochrome c oxidase biogenesis: Recent developments. Semin Cell Dev Biol. 2018;76:163–78. Epub 2017/09/06. doi: 10.1016/j.semcdb.2017.08.055. PubMed PMID: 28870773; PMCID: PMC5842095.

21. He J, Ford HC, Carroll J, Douglas C, Gonzales E, Ding S, Fearnley IM, Walker JE. Assembly of the membrane domain of ATP synthase in human mitochondria. Proc Natl Acad Sci U S A. 2018;115(12):2988–93. Epub 2018/02/15. doi: 10.1073/pnas.1722086115. PubMed PMID: 29440398; PMCID: PMC5866602.

22. Perez-Perez R, Lobo-Jarne T, Milenkovic D, Mourier A, Bratic A, Garcia-Bartolome A, Fernandez-Vizarra E, Cadenas S, Delmiro A, Garcia-Consuegra I, Arenas J, Martin MA, Larsson NG, Ugalde C. COX7A2L Is a Mitochondrial Complex III Binding Protein that Stabilizes the III2+IV Supercomplex without Affecting Respirasome Formation. Cell Rep. 2016;16(9):2387–98. Epub 2016/08/23. doi: 10.1016/j.celrep.2016.07.081. PubMed PMID: 27545886; PMCID: PMC5007171.

23. Lu L, Chen L, Gao F, Xu C, Ni C, Qian W. Rab5if is a potential therapeutic target of NSCLC. Cancer Genet. 2025;294-295:123–35. Epub 2025/05/04. doi: 10.1016/j.cancergen.2025.04.005. PubMed PMID: 40318299.

24. Liu X, Hajnoczky G. Altered fusion dynamics underlie unique morphological changes in mitochondria during hypoxia-reoxygenation stress. Cell Death Differ. 2011;18(10):1561–72. Epub 2011/03/05. doi: 10.1038/cdd.2011.13. PubMed PMID: 21372848; PMCID: PMC3172112.

25. Ahmad T, Aggarwal K, Pattnaik B, Mukherjee S, Sethi T, Tiwari BK, Kumar M, Micheal A, Mabalirajan U, Ghosh B, Sinha Roy S, Agrawal A. Computational classification of mitochondrial shapes reflects stress and redox state. Cell Death Dis. 2013;4(1):e461. Epub 2013/01/19. doi: 10.1038/cddis.2012.213. PubMed PMID: 23328668; PMCID: PMC3564000.

26. Bleck CKE, Kim Y, Willingham TB, Glancy B. Subcellular connectomic analyses of energy networks in striated muscle. Nat Commun. 2018;9(1):5111. Epub 2018/12/07. doi: 10.1038/s41467-018-07676-y. PubMed PMID: 30504768; PMCID: PMC6269443.

27. Sun Y, Daemen A, Hatzivassiliou G, Arnott D, Wilson C, Zhuang G, Gao M, Liu P, Boudreau A, Johnson L, Settleman J. Metabolic and transcriptional profiling reveals pyruvate dehydrogenase kinase 4 as a mediator of epithelial-mesenchymal transition and drug resistance in tumor cells. Cancer Metab. 2014;2(1):20. Epub 2014/11/08. doi: 10.1186/2049-3002-2-20. PubMed PMID: 25379179; PMCID: PMC4221711.

28. Liu X, Cooper DE, Cluntun AA, Warmoes MO, Zhao S, Reid MA, Liu J, Lund PJ, Lopes M, Garcia BA, Wellen KE, Kirsch DG, Locasale JW. Acetate Production from Glucose and Coupling to Mitochondrial Metabolism in Mammals. Cell. 2018;175(2):502–13 e13. Epub 2018/09/25. doi: 10.1016/j.cell.2018.08.040. PubMed PMID: 30245009; PMCID: PMC6173642.

29. Wang X, Xu B, Du J, Xia J, Lei G, Zhou C, Hu J, Zhang Y, Chen S, Shao F, Yang J, Li Y. Characterization of pyruvate metabolism and citric acid cycle patterns predicts response to immunotherapeutic and ferroptosis in gastric cancer. Cancer Cell Int. 2022;22(1):317. Epub 2022/10/14. doi: 10.1186/s12935-022-02739-z. PubMed PMID: 36229828; PMCID: PMC9563156.

30. Xia J, Li S, Liu S, Zhang L. Aldehyde dehydrogenase in solid tumors and other diseases: Potential biomarkers and therapeutic targets. MedComm (2020). 2023;4(1):e195. Epub 2023/01/26. doi: 10.1002/mco2.195. PubMed PMID: 36694633; PMCID: PMC9842923.

31. Martin MJ, Eberlein C, Taylor M, Ashton S, Robinson D, Cross D. Inhibition of oxidative phosphorylation suppresses the development of osimertinib resistance in a preclinical model of EGFR-driven lung adenocarcinoma. Oncotarget. 2016;7(52):86313–25. Epub 2016/11/20. doi: 10.18632/oncotarget.13388. PubMed PMID: 27861144; PMCID: PMC5349916.

32. Dobin A, Davis CA, Schlesinger F, Drenkow J, Zaleski C, Jha S, Batut P, Chaisson M, Gingeras TR. STAR: ultrafast universal RNA-seq aligner. Bioinformatics. 2013;29(1):15–21. Epub 2012/10/30. doi: 10.1093/bioinformatics/bts635. PubMed PMID: 23104886; PMCID: PMC3530905.

33. Liao Y, Smyth GK, Shi W. featureCounts: an efficient general purpose program for assigning sequence reads to genomic features. Bioinformatics. 2014;30(7):923–30. Epub 2013/11/15. doi: 10.1093/bioinformatics/btt656. PubMed PMID: 24227677.

34. Robinson MD, Oshlack A. A scaling normalization method for differential expression analysis of RNA-seq data. Genome Biol. 2010;11(3):R25. Epub 2010/03/04. doi: 10.1186/gb-2010-11-3-r25. PubMed PMID: 20196867; PMCID: PMC2864565.

35. Chen Y, Chen L, Lun ATL, Baldoni PL, Smyth GK. edgeR v4: powerful differential analysis of sequencing data with expanded functionality and improved support for small counts and larger datasets. Nucleic Acids Res. 2025;53(2). Epub 2025/01/23. doi: 10.1093/nar/gkaf018. PubMed PMID: 39844453; PMCID: PMC11754124.

36. Ritchie ME, Phipson B, Wu D, Hu Y, Law CW, Shi W, Smyth GK. limma powers differential expression analyses for RNA-sequencing and microarray studies. Nucleic Acids Res. 2015;43(7):e47. Epub 2015/01/22. doi: 10.1093/nar/gkv007. PubMed PMID: 25605792; PMCID: PMC4402510.

37. Kuleshov MV, Jones MR, Rouillard AD, Fernandez NF, Duan Q, Wang Z, Koplev S, Jenkins SL, Jagodnik KM, Lachmann A, McDermott MG, Monteiro CD, Gundersen GW, Ma’ayan A. Enrichr: a comprehensive gene set enrichment analysis web server 2016 update. Nucleic Acids Res. 2016;44(W1):W90-7. Epub 2016/05/05. doi: 10.1093/nar/gkw377. PubMed PMID: 27141961; PMCID: PMC4987924.

38. Chen EY, Tan CM, Kou Y, Duan Q, Wang Z, Meirelles GV, Clark NR, Ma’ayan A. Enrichr: interactive and collaborative HTML5 gene list enrichment analysis tool. BMC Bioinformatics. 2013;14:128. Epub 2013/04/17. doi: 10.1186/1471-2105-14-128. PubMed PMID: 23586463; PMCID: PMC3637064.

39. Xie Z, Bailey A, Kuleshov MV, Clarke DJB, Evangelista JE, Jenkins SL, Lachmann A, Wojciechowicz ML, Kropiwnicki E, Jagodnik KM, Jeon M, Ma’ayan A. Gene Set Knowledge Discovery with Enrichr. Curr Protoc. 2021;1(3):e90. Epub 2021/03/30. doi: 10.1002/cpz1.90. PubMed PMID: 33780170; PMCID: PMC8152575.

40. Chaudhry A, Shi R, Luciani DS. A pipeline for multidimensional confocal analysis of mitochondrial morphology, function, and dynamics in pancreatic beta-cells. Am J Physiol Endocrinol Metab. 2020;318(2):E87–E101. Epub 2019/12/18. doi: 10.1152/ajpendo.00457.2019. PubMed PMID: 31846372; PMCID: PMC7052579.

41. Li H. New strategies to improve minimap2 alignment accuracy. Bioinformatics. 2021;37(23):4572–4. Epub 2021/10/09. doi: 10.1093/bioinformatics/btab705. PubMed PMID: 34623391; PMCID: PMC8652018.

42. Weissensteiner H, Forer L, Fendt L, Kheirkhah A, Salas A, Kronenberg F, Schoenherr S. Contamination detection in sequencing studies using the mitochondrial phylogeny. Genome Res. 2021;31(2):309–16. Epub 2021/01/17. doi: 10.1101/gr.256545.119. PubMed PMID: 33452015; PMCID: PMC7849411.

43. Rath S, Sharma R, Gupta R, Ast T, Chan C, Durham TJ, Goodman RP, Grabarek Z, Haas ME, Hung WHW, Joshi PR, Jourdain AA, Kim SH, Kotrys AV, Lam SS, McCoy JG, Meisel JD, Miranda M, Panda A, Patgiri A, Rogers R, Sadre S, Shah H, Skinner OS, To TL, Walker MA, Wang H, Ward PS, Wengrod J, Yuan CC, Calvo SE, Mootha VK. MitoCarta3.0: an updated mitochondrial proteome now with sub-organelle localization and pathway annotations. Nucleic Acids Res. 2021;49(D1):D1541-D7. Epub 2020/11/12. doi: 10.1093/nar/gkaa1011. PubMed PMID: 33174596; PMCID: PMC7778944.

44. Auwera GAVd, O’Connor BD. Genomics in the Cloud : Using Docker, GATK, and WDL in Terra. Sebastopol: O’Reilly Media, Incorporated; 2020. Available from: https://search.ebscohost.com/login.aspx?direct=true&scope=site&db=nlebk&db=nlabk&AN=2420556.

